# Further Perceptions of Probability: In Defence of Associative Models – A Commentary on Gallistel et al. 2014

**DOI:** 10.1101/2020.01.30.927558

**Authors:** Mattias Forsgren, Peter Juslin, Ronald van den Berg

## Abstract

Extensive research in the behavioural sciences has addressed people’s ability to learn stationary probabilities, which stay constant over time, but only recently have there been attempts to model the cognitive processes whereby people learn – and track – *non-stationary* probabilities. In this context, the old debate on whether learning occurs by gradual formation of associations or by occasional shifts between hypotheses representing beliefs about distal states of the world has resurfaced. Gallistel et al. (2014) pitched the two theories against each other in a non-stationary probability learning task. They concluded that various qualitative patterns in their data were incompatible with trial-by-trial associative learning and could only be explained by a hypothesis-testing model. Here, we contest that claim and demonstrate that it was premature. First, we argue that their experimental paradigm consisted of two distinct tasks: probability tracking (an estimation task) and change detection (a decision-making task). Next, we present a model that uses the (associative) delta learning rule for the probability tracking task and bounded evidence accumulation for the change-detection task. We find that this combination of two highly established theories accounts well for all qualitative phenomena and outperforms the alternative model proposed by Gallistel et al. in a quantitative model comparison. In the spirit of cumulative science, we conclude that current experimental data on human learning of non-stationary probabilities can be explained as a *combination* of associative learning and bounded evidence accumulation and does not require a new model.

## INTRODUCTION

The issue of how people learn and assess probabilities has been pivotal to the behavioural sciences at least since the Enlightenment and studied extensively, especially in psychology and behavioural economics. Typically, this has occurred in the context of assuming *stationary probabilities* in the environment (i.e., probabilities that stay constant over time). This research shows that people are good at learning probabilities from experience with relative frequencies (Edwards, 1961; Estes, 1976; Fiedler, 2000; Peterson & Beach, 1967). Yet research on heuristics and biases shows that probability assessments are sometimes swayed by subjective (“intentional”) aspects, like prototype-similarity (representativeness) or ease of retrieval, leading to biased judgements (Kahneman & Frederick, 2005). People also appear to over-weight extreme probabilities in their decisions when encountering them in numeric form (Tversky & Kahneman, 1992), but under-weight them when they are learned inductively from trial-by-trial experience (Hertwig & Erev, 2009). People frequently have problems with reasoning according to probability theory, leading to phenomena like base-rate neglect and conjunction fallacies (Kahneman & Frederick, 2005; Tversky & Kahneman, 1983), at least if they cannot benefit from natural frequency formats (Gigerenzer & Hoffrage, 1995) that highlight the set-relations between the events (Barbey & Sloman, 2007).

Not all probabilities are stationary, as when, for example, the risks of default in a mortgage market fluctuate over time or the risk of hurricanes changes with a changing global climate. Since modelling how humans learn – and track – *non-stationary probabilities* involves changes in people’s beliefs about probability, it has (once again) highlighted the classical issue of whether people learn gradually, trial-by-trial, by forming *associations* between concepts or by occasional shifts between discrete *hypotheses* about the state of the world (Bruner, Goodnow, & Austin, 1956). These two kinds of models have different historical origins and – in their original formulations – make quite different claims about the learning process. Associative models in the empiricist tradition treat learning as a process in which associations are gradually formed between contiguous stimuli. Importantly, this kind of learning does not involve propositions about the world that can be true or false. Hypothesis testing models in the rationalist tradition, by contrast, treat learning as a process of forming propositional beliefs about (potentially unobservable) distal states of the world, which by definition are either true or false.

Although this orthodox epistemic interpretation of the claims of the models is less common now, there still exists a plethora of models in cognitive science and neuroscience that in various ways refer to “associative trial-by-trial learning” or to the “testing and updating of hypotheses”. A neuropsychological and psychophysical literature suggests a cohort of probability learning models that estimate in a trial-by-trial manner (Nassar et al., 2012; Nassar, Wilson, Heasly, & Gold, 2010; Norton, Acerbi, Ma, & Landy, 2019; Wilson, Nassar, & Gold, 2013; Wilson, Nassar, Tavoni, & Gold, 2018) which is supported by findings that learning rates are modulated by trial-level prediction errors registered in the anterior cingulate cortex (Behrens, Woolrich, Walton, & Rushworth, 2007; Rushworth & Behrens, 2008; Silvetti, Seurinck, & Verguts, 2013). However, these ideas have been challenged by a small, mostly recent literature (Gallistel, Krishan, Liu, Miller, & Latham, 2014; Khaw, Stevens, & Woodford, 2017; Ricci & Gallistel, 2017; Robinson, 1964). In particular, empirical phenomena related to trial-by-trial probability estimates and explicit reports of changes in the underlying probability have been claimed (Gallistel et al., 2014; Ricci & Gallistel, 2017) to be incompatible with associative trial-by-trial models and to require a model built on hypothesis testing.

In this Theoretical Note, we challenge these claims. We propose that construing the learning of non-stationary probabilities as *either* based on associative models *or* hypothesis testing models is not a fruitful way to frame the question.^1^ The experimental paradigm which has led researchers to support a hypothesis-testing model consists, we argue, of two distinct tasks. The first one is to track the frequency of observed stimuli – the traditional domain of associationism. The second one is to detect changes in the process that generates the stimuli, which involves the testing of discrete hypotheses about hidden states of the world – a decision-making task about propositional beliefs that is well suited to a rationalist approach. Given the highly different natures of these two tasks, we propose that they may be driven by distinct mechanisms. To test this, we construct a model that uses delta-rule learning for estimating the probability and bounded evidence accumulation for the change detection task. To preview our results, we find that this combination of two highly established models accounts well for all reported phenomena and outperforms the purely rationalist model proposed by Gallistel et al. (2014) in a quantitative model comparison on data from three previous studies. Hence, the claims by Gallistel et al. were premature.

### Tracking Probabilities in Non-Stationary Environments

While there is a large literature on how people learn stationary probabilities, there are only a few studies that have addressed learning of non-stationary probabilities. In the studies revisited here, participants were asked to estimate the hidden Bernoulli parameter by adjusting a physical lever (Robinson, 1964) or a slider on a computer screen (Gallistel et al., 2014; Khaw et al., 2017; Ricci & Gallistel, 2017). In the latter case, this was framed as the proportion of green rings in a hypothetical box visualised on a computer screen (Figure 1A). On each trial, the participant could adjust a slider between 0 and 100 percent to produce an estimate, before locking in their guess and initiating the next draw from the box (i.e., the next trial). The participants performed 10,000 trials and, importantly, were free to choose to revise their estimate or to leave it unchanged on any trial. The data of interest are the realised outcomes from the Bernoulli process, the underlying true probabilities of the outcomes, and the participant’s estimates of these underlying probabilities (Figure 1B). Most participants exhibited stepwise updating behaviour: for long periods they did not adjust their estimates, at other times more often, but never on every trial. One of the studies (Gallistel et al., 2014) included a button labelled “I think the box has changed” that allowed participants to indicate that they believed that there had been a change in the parameter of the Bernoulli process. Half of those participants were also provided with the option to retract their decisions by pressing a button labelled “I take that back” (so called “second thoughts”, see Figure 1A).

**Figure 1.**
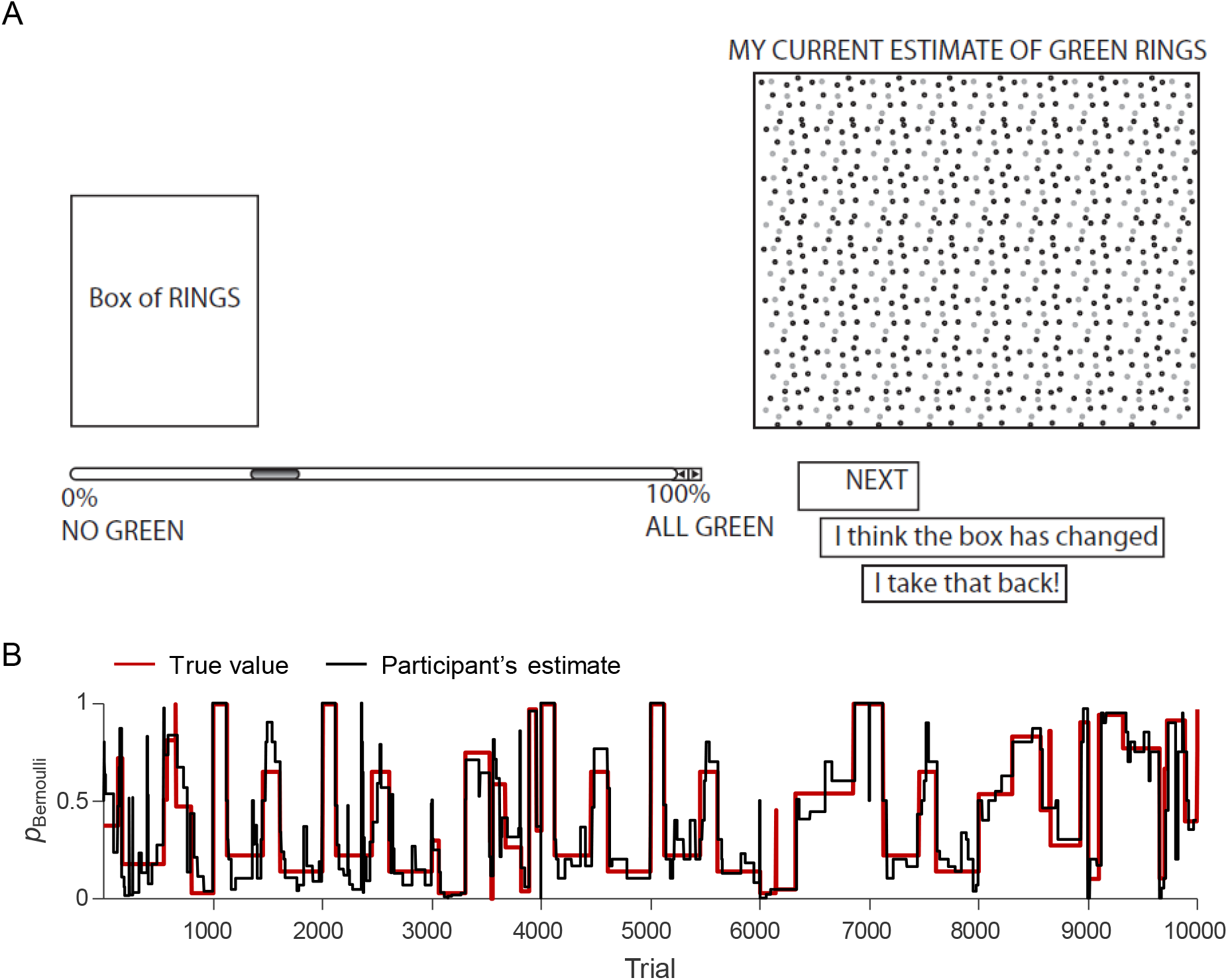
Experimental paradigm. (A) Screenshot of the task in Gallistel et al (2014). Khaw, Stevens and Woodford (2017) and Ricci and Gallistel (2017) used a similar design but without buttons for reporting that the box has changed or second thoughts. From “The perception of probability,” by C. R. Gallistel, M. Krishan, Y. Liu, R. Miller and P. E. Latham, 2014, *Psychological review, Vol. 121*, p. 96-123. Copyright 2014 by American Psychological Association. Reprinted with permission. (B) Example of response data (black) in an experiment where the hidden Bernoulli probability (red) was non-stationary and stepwise (Participant 1 in Gallistel et al, 2014).

### Two Approaches to Learning: Forming Associations *vs*. Testing Hypotheses

As in many areas of the psychology of learning, there are two different ways of explaining how people infer probabilities from experience. Models with their origin in the associationist traditions of behaviourism, reinforcement learning, and connectionism emphasise the continuous updating of associations “trial-by-trial”, while models with their origin in cognitive psychology often emphasise the testing of and discrete shifting between hypotheses.

The most famous associationist theory is perhaps the Rescorla-Wagner model of classical conditioning (Rescorla & Wagner, 1972), which has been adopted in many other domains too (Behrens et al., 2007; Busemeyer & Myung, 1988; Neal & Dayan, 1997; Verguts & Van Opstal, 2014). This model is based on the delta learning rule introduced by Widrow and Hoff (1960), an algorithm for updating the weights of nodes in a neural network (see Widrow & Lehr, 1993, for a review). A defining feature of the delta rule is that it is an “eager” algorithm: the associations are updated each time a new data point is observed (“trial-by-trial updating”). For the task of learning a binomial probability from observed outcomes, the delta-rule can be implemented as

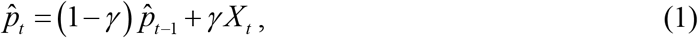

where 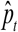 is the probability estimate at time *t*, 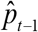 the previous estimate, *γ* the learning rate, and *X_t_* ∈{0,1} the observed outcome (e.g., “0 = red”, “1 = green”). The delta-rule accordingly abstracts an online running estimate of the underlying probability in the form of an associative expectation of the event. It has the advantage of being recursive: it operates without requiring access to memories going back further than the latest observation.

By contrast, models in the rationalist tradition typically assume that people construct richer representations of reality than mere associations between contiguous features, instead referring to beliefs about distal states of the world that can be either true or false. One way to embody this is in hypothesis-testing models. These assume that people learn about the world by testing between explicit hypotheses about the state of the world based on confirming or disconfirming feedback (Brehmer, 1974; Bruner et al., 1956). A common feature of such models is that beliefs are updated in a discrete rather than gradual fashion, because observers hold on to a belief until sufficiently strong evidence has accumulated against it. Hypothesis testing models have been applied to, for example, research on reasoning (e.g. Klayman & Ha, 1987; Oaksford & Chater, 1994; Wason & Johnson-Laird, 1970), categorisation (Ashby & Valentin, 2017; Bruner et al., 1956), and function learning (Brehmer, 1974, 1980). Because a single data point typically provides little evidence about a hypothesis, these models predict that beliefs sometimes stay unchanged over many outcome observations. Gallistel et al. (2014) formalised a hypothesis-testing model for the learning of non-stationary probabilities which they called the “If it ain’t broke, don’t fix it” (IIAB) model. According to this model, participants assess after each new observation whether their current belief is “broke” and only update it if they decide that it is. Thus, humans do not learn probabilities directly: they learn “change points” in the hidden Bernoulli parameter and use this information and memories of the previous outcomes to *infer* probabilities.

### Empirical Phenomena Related to Human Estimation of Non-Stationary Probabilities

To evaluate the plausibility of these two classes of models in the context of non-stationary probability learning, Gallistel et al. (2014) identified a number of important empirical phenomena that any viable model should be able to reproduce. Table 1 provides an overview of these phenomena, which we divide into two categories: those related directly to slider updates and those related to participants’ stated beliefs about the generative function.

**Table 1.**
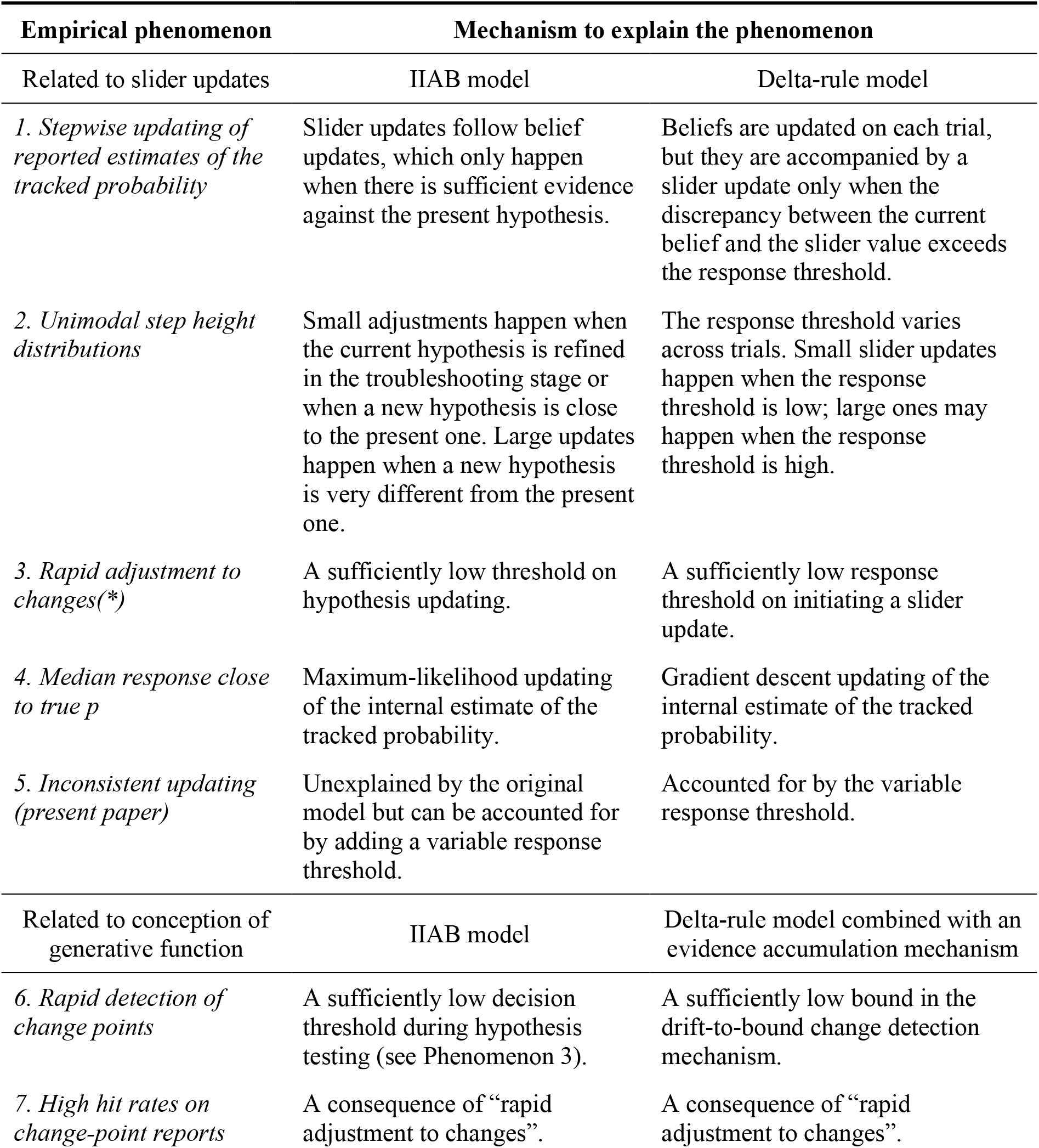
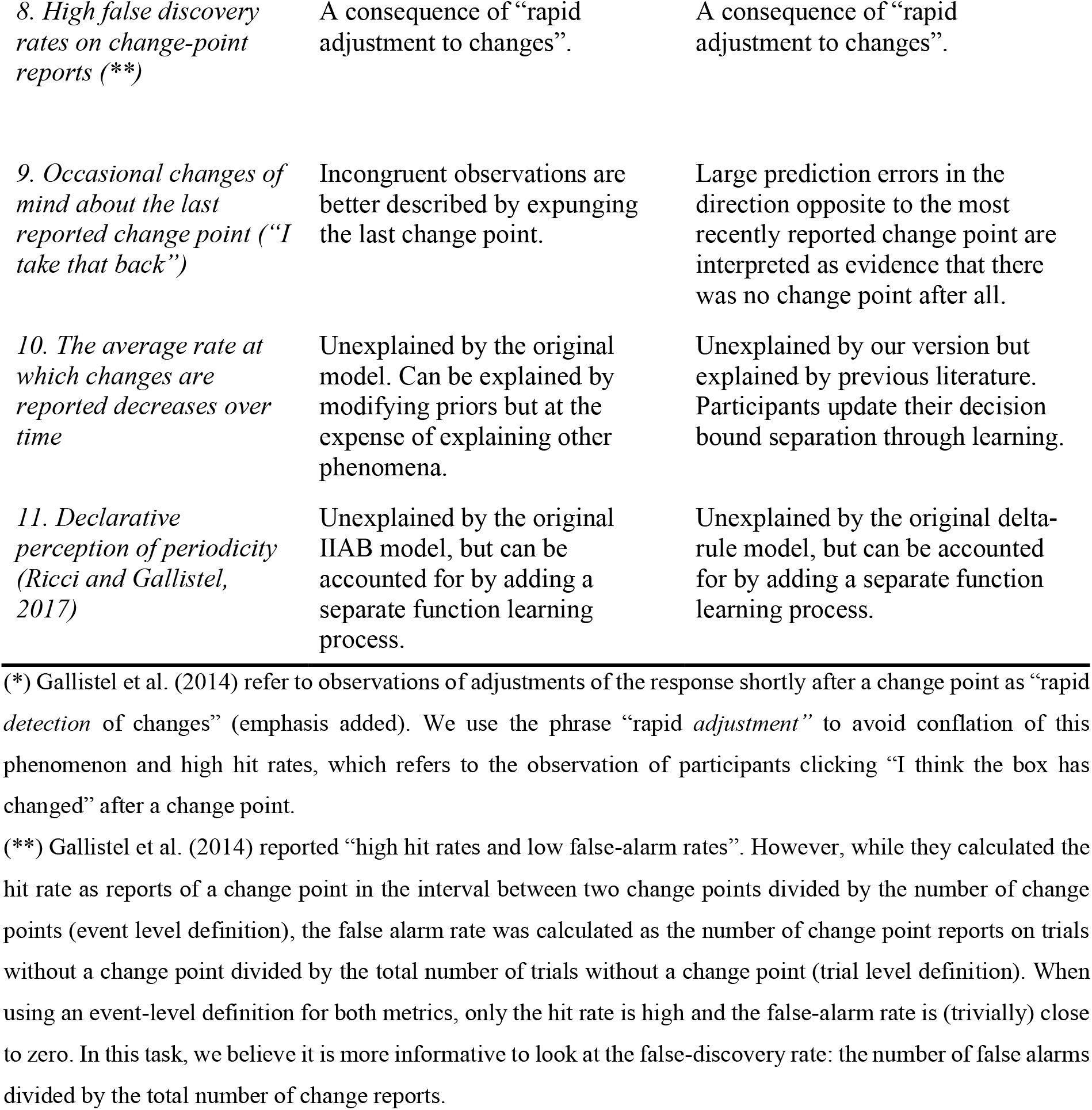
Empirical Phenomena Observed in the Probability Tracking Task with the Mechanisms of the IIAB and the Proposed Mechanisms of our Extended Delta-rule and Evidence Accumulation model that Explain them. All Phenomena Reported by Gallistel et al. (2014) Unless Indicated Otherwise.

We identified an additional phenomenon that has not been reported before but may be informative about the underlying mechanisms: participants regularly make changes to the slider in the *opposite* direction of the colour of the last observation (e.g., decrease their estimate of the probability of a red outcome after observing a red outcome). In the three datasets considered in the present study, 23.6±1.6% (M±SE across participants) of the updates were of this nature. This phenomenon is unexplainable by both the IIAB model and the standard delta-rule model, but we will show how both models can account for it by extending them with a variable response threshold.

There are two more phenomena (10 & 11) that cannot be explained by either of the models, at least not as specified here. We believe both models could explain them with certain extensions, which we will come back to in the Discussion. Because these phenomena are a shared issue, and thus not diagnostic of the learning mechanisms that we are contrasting here, we will not consider them in our evaluations of the models.

### The Main Arguments Against Associative Models

Gallistel et al. (2014) argue that associative models, which update an internal estimate trial-by-trial, are unable to account for several of the phenomena listed above. The first one is the stepwise manner in which participants tend to adjust their estimates of tracked probabilities: they often leave the slider unchanged for long periods of time (Figure 1B), which seems in direct contradiction to any model that updates on a trial-by-trial basis. An additional and closely related argument against those models is based on the distribution of adjustment sizes. Besides making many small adjustments, participants also regularly make large adjustments. Large adjustments are hard to reconcile with the idea of a gradual build-up of an associative value because a single observation should rarely cause a large change in that estimate. Gallistel et al. (2014) argue that large adjustments and periods of constancy instead reflect discrete belief changes as the participant moves from conclusion to conclusion.

One potential way to make an associationist model account both for large slider adjustments and periods with no adjustment is to assume that participants have a “response threshold” that prevents them from making slider updates when the difference between the current slider value and their internal belief is not sufficiently large to justify the effort. Such a threshold could reflect simple “laziness” (recall that participants typically performed thousands of trials) or have a more sophisticated basis. While a response threshold produces stepwise response behaviour, it has another problem: it is unable to explain adjustments smaller than the threshold. As noted by Gallistel et al. (2014), a potential remedy is to make one further assumption, namely that the response threshold can vary across trials. This variability could reflect, for example, fluctuations in attention and motivation, or noise in neural and cognitive processing (Drugowitsch, Wyart, Devauchelle, & Koechlin, 2016; Faisal, Selen, & Wolpert, 2008). A more sophisticated proposal is that the variable threshold reflects a rational process in which participants trade off costs related to moving the mouse and cognitive processing against accuracy in task performance (Khaw et al., 2017). Gallistel et al. (2014) inspected the behaviour of a trial-by-trial model with a variable response threshold through simulations but were unable to find parameter settings that produced step height distributions resembling the empirically observed distributions. Importantly, however, they seem to have done this by manually trying out different parameter settings, rather than by exploring the space exhaustively. In the present paper, we perform a more systematic search and find that a delta-rule model with a variable response threshold does, in fact, accurately reproduce the empirical distributions.

Another major argument that Gallistel et al. (2014) make against associative models is based on their observation that participants are able to detect changes in the Bernoulli parameter (Phenomena 6-9 in Table 1). They demonstrated this using a version of the task where participants were asked to press an “I think the box has changed” button whenever they thought that there had been a change in the generative process (see Figure 1A). Some of these participants were also given the option to report “seconds thoughts” about those reports by pushing a button labelled “I take that back.” (Figure 1A). Gallistel et al. (2014) interpret the ability to detect and reconsider changes as evidence that participants store a record of the previous change points in memory, over and above a summary representation of the outcomes thus far observed. They argue that such a record is incompatible with associative models, which have a more condensed knowledge state. In the present work, we argue that estimating a probability and detecting changes in the underlying process are two separate tasks that may be driven by separate mechanisms. We show that a delta-rule model extended with a bounded evidence accumulation mechanism tracks changes in the underlying Bernoulli parameter and can account for human reports of, and second thoughts about, such changes.

Based on the above arguments, Gallistel et al. (2014) ruled out the entire class of associative, trial-by-trial estimation models as a possible explanation for human behaviour on probability estimation tasks. They argued that one instead needs a model with the conceptual richness of a hypothesis testing, “troubleshooting” process that identifies the most likely state of the world to have produced the data. They proposed such a model under the name “If It Ain’t Broke don’t fix it” (IIAB, described in more detail later) and used simulations to show that there are parameter settings that produce qualitatively similar data patterns as the phenomena observed in the human data. Importantly, however, they did not fit the model to any data and they did not perform any quantitative model comparison against alternative models.

### Outline of this paper

Gallistel et al. (2014) argued that associative models (represented by the delta-rule) are qualitatively incompatible with human estimation of non-stationary probabilities. We present a re-evaluation of that claim and an extension of their analyses. In the next section, we present the two main contending models: the IIAB hypothesis-testing model proposed by Gallistel et al. (2014) and a trial-by-trial estimation model based on delta-rule learning. Thereafter, we use simulations to examine if the delta-rule model can reproduce the previously emphasised qualitative aspects of human data related to the slider updates (Phenomena 1 to 5 in Table 1). Unlike Gallistel et al. (2014), we find that it accurately reproduces those patterns.

We further note that to account for all of the phenomena in Table 1, a model may need two separate mechanisms: one for tracking the proximal stimulus distribution (the recently observed frequencies) and another one that uses this information to make decisions about the likely distal state of the world (the underlying generative probability). We demonstrate that combining the delta-rule’s associative account of the descriptive tracking of the probability with a sequential evidence accumulator on the delta-rule’s prediction error is sufficient to account for the explicitly stated inferences about the generative process.

Having established that there are no qualitative reasons to rule out the delta-rule model, we next examine how well both models account for actual data by fitting them to data from the three previous studies. We find that both models account well for most of the data, even though formal model comparison based on cross validation clearly favours the associative model over the IIAB model for almost every participant. To paraphrase Mark Twain (White, 1897), our results indicate that the report of the death of associative models was an exaggeration.

## MODELLING

### The IIAB Model

We provide a brief description of the “If It Ain’t Broke, don’t fix it” (IIAB) model here and refer the reader to Gallistel et al. (2014) for a more complete exposition and mathematical details. A key characteristic of this model is that it has a relatively stable internal belief about the tracked probability^2^: it only updates this belief when there is sufficient evidence against the current value. It proceeds in two stages. In the first stage, it tests whether the currently held belief about the tracked probability is “broke”. This test is performed by computing the discrepancy between the belief and the outcomes observed since the last registered change point. If the discrepancy – measured as Kullback-Leibler divergence – exceeds a decision threshold *T*_1_, it is concluded that something is “broke”. Each time this happens, the model enters a “troubleshooting” stage, in which it considers three hypotheses on why the current estimate may be “broke”: (i) there was a change in the generative process (“I think the box has changed”), in which case the model will register a new change point and update its estimate of *p*_true_ accordingly; (ii) the previously registered change point was a mistake (“I take that back”), in which case the model will expunge the last recorded change point and update its estimate accordingly; (iii) the previous estimate of *p*_true_ was wrong but the change point record is correct, in which case the model will update its estimate of *p*_true_ but not register or expunge any change point. Hypothesis (iii) corresponds to concluding that the estimate was “broke” due to sampling error, but it is not assumed that such beliefs are recorded in memory. Gallistel et al. (2014) argue that these “troubleshooting” steps allow the IIAB model to explain behavioural phenomena related to the participant’s knowledge about the generative function (Phenomena 6-11 in Table 1). Because the updated estimate is always the average of all observations since the last change point and the model can have “second thoughts” about that change point, the model must keep a descriptive online tracking of the recent environmental input in addition to its estimate of *p*_true_. This description consists of a proportion, a running Kullback-Leibler divergence, and a vector of all observations after the second to last change-point.

The original version of the IIAB model has just two parameters: threshold *T*_1_ mentioned above and an additional threshold *T*_2_ that is used in the troubleshooting stage. While both thresholds are fixed, the evidence in the first stage is scaled by the number of trials since the last change (the sample size). The IIAB model will therefore become increasingly sensitive to small discrepancies between the current belief and the most recently observed evidence when no change point has been detected for a while.

Predictions related to slider updates can be derived directly from the model’s internal belief state about the tracked probability. Predictions related to a participant’s reports of suspected changes in the generative process and changes of mind about those reports can be derived directly from the model’s “troubleshooting” stage. Hence, this is a rich model that makes predictions about all of the empirical phenomena listed in Table 1 except 11.

### The Delta-rule Model

Delta-rule models update an estimate of the observed frequency after *every* observation. The most basic version does this using a recursive function of the form specified in Eq. (1) above. The model proceeds by constantly adjusting its estimate of the tracked probability in the direction of the latest observed outcome: seeing a green ring slightly increases the observer’s estimate of the proportion of green rings in the box and seeing a red one decreases it.

The higher the value of the learning rate, *λ*, the larger the trial-by-trial adjustments. In environments with frequent, abrupt changes in the generative process, it is beneficial to have a high learning rate because that will allow the model to catch up quickly to those changes. However, in stable or very slowly changing environments it is better to have a slow learning rate, to avoid the estimates being overly sensitive to occasional unexpected outcomes. The tasks used in previous studies on non-stationary probability tracking (Gallistel et al., 2014; Khaw et al., 2017; Nassar et al., 2010; Norton et al., 2019) are often a mixture of those two situations: long periods of stability with occasional, abrupt changes (Figure 1B). In such tasks, it can be a disadvantage to have a single, fixed learning rate.

Several modifications to the standard delta-rule model have been proposed that might work better in mixed environments, for example the addition of a second kernel (Gallistel et al., 2014) and the use of a dynamic learning rate (Nassar et al., 2010). However, it has been shown in a similar task that the basic model typically performs as well as or even better than more complex alternatives (Norton et al., 2019). While we will consider two variants later (see Results), our main analyses will use the most basic, single-parameter version of the delta-rule, as specified by Equation (1).

### Cumulative Prediction Error and Changes in the Generative Process

Gallistel et al. (2014) rightly point out that the delta-rule by itself cannot account for participant data related to explicit change point reports (Phenomena 6-9 in Table 1). This is not surprising since the delta rule is a learning mechanism for proximal stimuli. To explain change point reports, it needs to be combined with a decision-making mechanism which discriminates between hypotheses about distal states of the world. One of the most established mechanisms for perceptual decision-making to date is that of bounded evidence accumulation (Bogacz et al., 2006; Ditterich, 2006; Ratcliff, 1978), which finds broad support in behavioural, neurophysiological, and computational studies (Ratcliff, 1978; Ratcliff, Smith, Brown, & McKoon, 2016; Wagenmakers, 2009). Evidence accumulators proceed by integrating evidence over time until a bound is crossed, at which point a decision is made.

In the present task, the trial-by-trial prediction error, 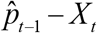, is indicative of changes in the tracked probability (Figure 2) and could thus be a suitable input variable for an evidence accumulator. When the generative process is stable and the observer’s estimate has homed in on a value close to the true value of the tracked variable, prediction errors tend to cancel each other out over trials and the cumulative evidence will hover around 0 (Figure 2, first 100 trials). After an abrupt change in the generative process (Figure 2, trial 100), however, there will typically be a burst of prediction errors that all have the same sign (determined by the direction of the change) and thus do not cancel each other out. Hence, the cumulative prediction error is indicative of changes in the generative process: a value close to zero suggests a stable process; a large positive value suggests that there was a recent increase in the Bernoulli parameter; a large negative value suggests that there was a recent decrease in the Bernoulli parameter. We propose that observers may use the cumulative prediction error to detect changes in the generative process when tasked to do so. We model this by adding a bounded evidence accumulator to the delta-rule model and let it trigger an “I think the box has changed” response whenever the cumulative prediction error exceeds a decision bound (Figure 2). A nice feature of this combination is that the evidence accumulator has the same computational efficiency as the delta-rule: the cumulative error can be updated iteratively with negligible memory requirements.

**Figure 2.**
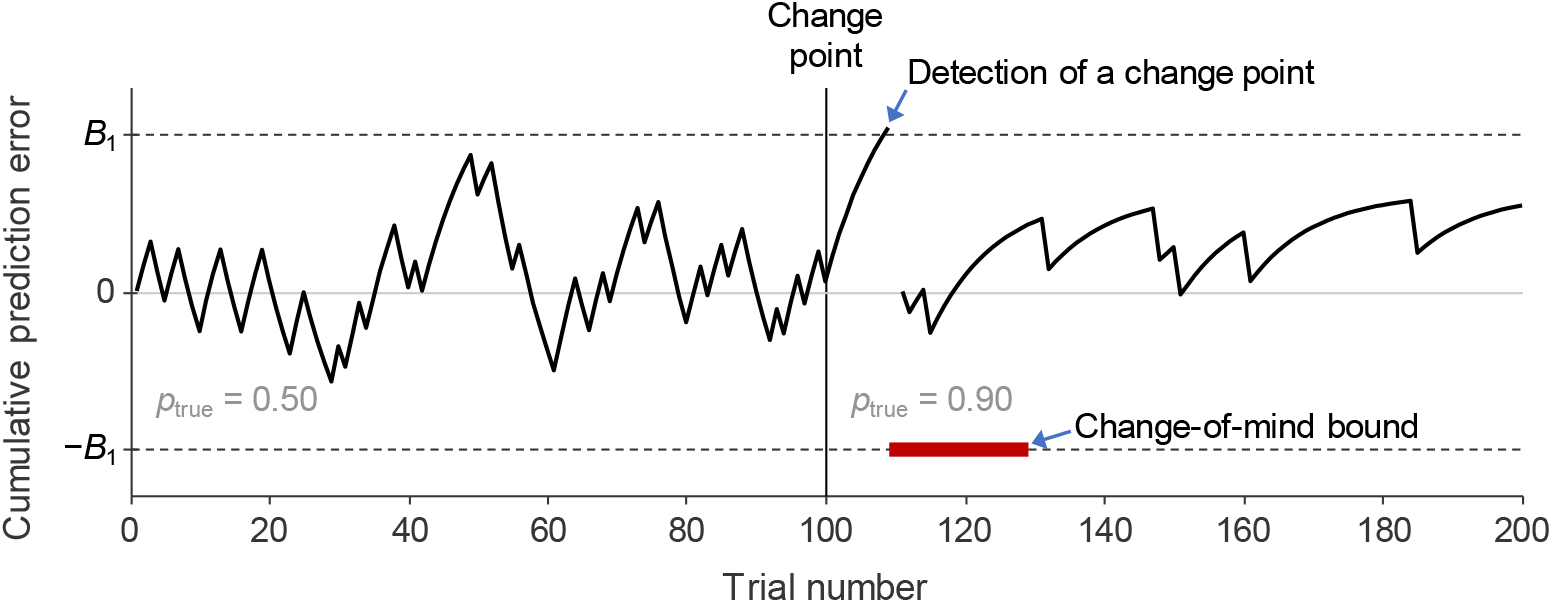
A proposed mechanism to detect changes in a Bernoulli process based on accumulation of prediction errors. Simulation of the cumulative prediction error in a delta-rule model with a learning rate of 0.90. The observer’s belief and the true value of the Bernoulli parameter, *p*_true_, are both initialized at 0.50. For the first 100 trials, the cumulative prediction error hovers around 0, because positive and negative errors cancel each other out (however, note that around trial 50 there is almost a false alarm). At trial 100, the value of *p*_true_ is changed to 0.90. This causes the cumulative prediction error to quickly increase, because now more positive errors are experienced than negative ones. The cumulative prediction error crosses decision bound *B*_1_=3.0 at trial 109 which triggers an “I think the box has changed” response, resets the cumulative prediction error to 0, and instates a temporary change-of-mind bound (which is not being crossed in this example). The reason why the shape of the cumulative prediction error looks different after the change in *p*_true_ is that the trial-by-trial prediction errors changed from being approximately −0.50 and 0.50 (in 50% of the trials each) before the change to around 0.90 (on 90% of the trials) and −0.10 (on 10% of the trials) after it.

Importantly, bounded evidence accumulators can also explain “second thoughts”, which are known as “changes of mind” in the decision-making literature. This is done by introducing a temporary second bound at the moment that an initial decision has been made (e.g., Resulaj, Kiani, Wolpert, & Shadlen, 2009; Van den Berg et al., 2016). This bound will be crossed if the immediate post-decision information is sufficiently inconsistent with the original decision, triggering a change-of-mind response. A typical way to implement this bound is to use two parameters, specifying its height and lifetime. Because we have very little data on changes of mind (115 reports by 5 participants in a total of 50,000 trials), we take a simpler approach by setting the change-of-mind bound equal to the original bound but in the opposite direction of the detected change point, such that the lifetime of the bound is the only additional parameter required to model these rare responses.

### Response Threshold

Previous studies (Gallistel et al., 2014; Khaw et al., 2017; Robinson, 1964) have considered the possibility that participants do not adjust the slider when the difference to their internal belief is too small. This could arise from participants economising their time costs^3^. Additionally, cognitive processes are noisy (Drugowitsch et al., 2016; Faisal et al., 2008) and participants’ levels of motivation and attention might fluctuate over time, which could make the discrepancy required for an update vary across trials. We will first model this resistance to updating in the same was as in Gallistel et al. (2014), namely by using a response threshold whose value is drawn on each trial from a constrained Gaussian distribution. We will then test models where the threshold is drawn from a beta distribution. We parameterise both thresholds by their mean and variance.

### Response Noise and Lapse Rate

To account for inaccuracies in predicted slider settings – due to factors such as motor noise and model mismatch – we included response noise in all models. This noise was implemented as a beta distribution centred on the model’s predicted response, *m*, and was applied to trials on which a slider update was predicted. Since the variance of the beta distribution has an upper bound (equal to *m* – *m*^2^), we parameterised it as a relative value between 0 (no variance) and 1 (maximum variance). The (relative) variance was fitted as a free parameter. Moreover, we included a small lapse rate (1/1000) to account for lapses in attention and to avoid numerical instabilities in model variants without any other sources of stochasticity (such as the original IIAB model). We performed several robustness checks (reported in Results) to verify that the results do not critically depend on these auxiliary assumptions.

### Model Evaluation Methods

We will evaluate the models by using maximum-likelihood fitting with five-fold cross-validation (see Supplemental Materials for details about the likelihood function and optimisation method). To verify that the models make clearly distinguishable predictions, and that our model comparison methods are able to detect these differences, we performed a model recovery analysis. We find that both model selection and parameter estimation are accurate (see Supplemental materials for details).

### Open science notes

Modelling code and the data can be found at https://osf.io/zhv2r/ (Forsgren et al., 2022). This study was not preregistered.

## RESULTS

In this section, we evaluate the models in two different ways. First, we perform simulations to re-assess Gallistel et al.’s (2014) conclusion that a delta-rule model cannot reproduce the main qualitative phenomena observed in human data (Table 1). Second, we use likelihood-based five-fold cross validation to compare this model with its main contender, the IIAB model.

### Reassessment of the Conclusion that Delta-rule Model Predictions are Qualitatively Inconsistent with Data

Gallistel et al. (2014) were unable to find parameter settings for the delta-rule model that produced distributions of step widths and step heights with similar shapes as observed in human data (Phenomena 1-3 in Table 1). From this, they concluded that delta-rule models are fundamentally unfit to account for human estimation of non-stationary probabilities. Here, we reconsider this finding by using an approach that differs from theirs in an important way: instead of manually trying out parameter settings, we systematically explore parameter space using an optimisation method. Specifically, we let the algorithm search for the setting that minimises the root mean squared deviation (RMSD) between the data and the model prediction for the summary statistic of interest (histograms of step width and height, cumulative number of updates, etc).

We use the exact same delta-rule model as tested by Gallistel et al. (2014), which has three parameters: the learning rate (*λ*), the mean of the (Gaussian) response threshold distribution (*μ*_τ_), and the coefficient of variation of this distribution (*cv*_τ_) (hence, no response noise or lapse rate was included in the model at this stage of the analysis). Just like Gallistel et al. (2014), we constrain *cv*_τ_ to have a maximum value of 0.33. In contrast to their findings, we find that this model reproduces the step width and step height distributions very well (Figure 3). It also does an excellent job in reproducing the other phenomena related to slider updates: the cumulative number of updates, the median response values, and the cumulative distribution of the latency between changes in the Bernoulli parameter and the next slider update.

**Figure 3.**
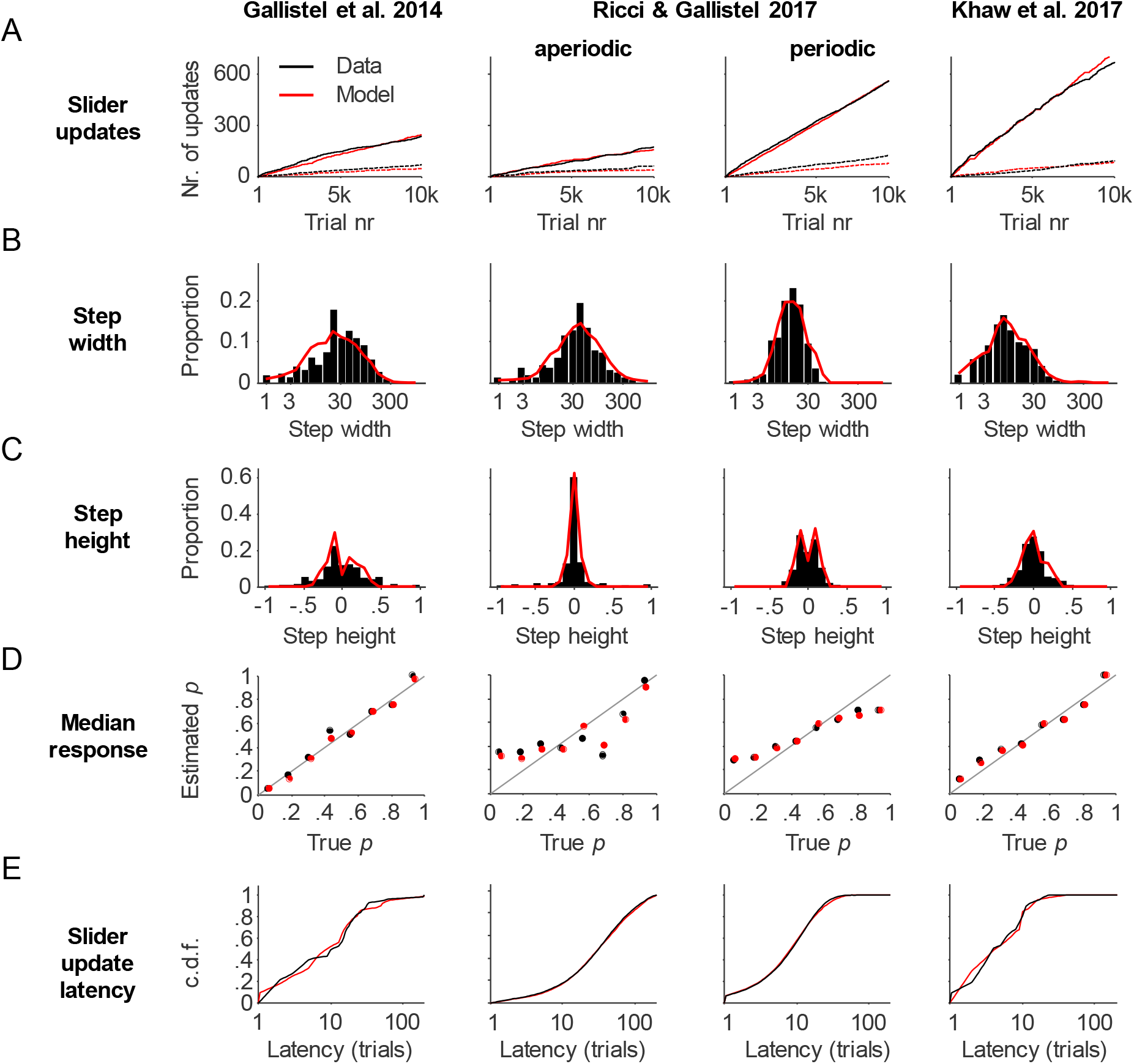
Evaluation of qualitative predictions by the delta-rule model related to slider settings. Results are shown for Participant 1 in each of the 4 analyzed datasets. The model simulations results (red) were obtained by minimizing the root mean squared deviation (RMSD) with the data (black). (A) Total number of slider updates (solid) and number of inconsistent slider updates (dashed) as a function of trial number. (B) Distribution of the number of trials between consecutive slider updates. (C) Distribution of the magnitude of slider updates on trials with an update. (D) Median estimate of the tracked probability versus the median true value. (E) Cumulative distribution of the number of trials between a change in *p*_true_ and the next slider update.

Next, we extend the model with sequential accumulation of the prediction error and test if the resulting model can account for phenomena related to the conception of the generative function (Phenomena 6-9 in Table 1). We find that the model accurately reproduces these phenomena too (Figure 4): the cumulative number of “I think the box has changed” responses; the cumulative number of “I take that back” responses; the cumulative distribution of the latency between a change in the generative function and the observer’s detection of the change; the hit rates, false discovery rates, and false alarm rates of box-change detections.

**Figure 4.**
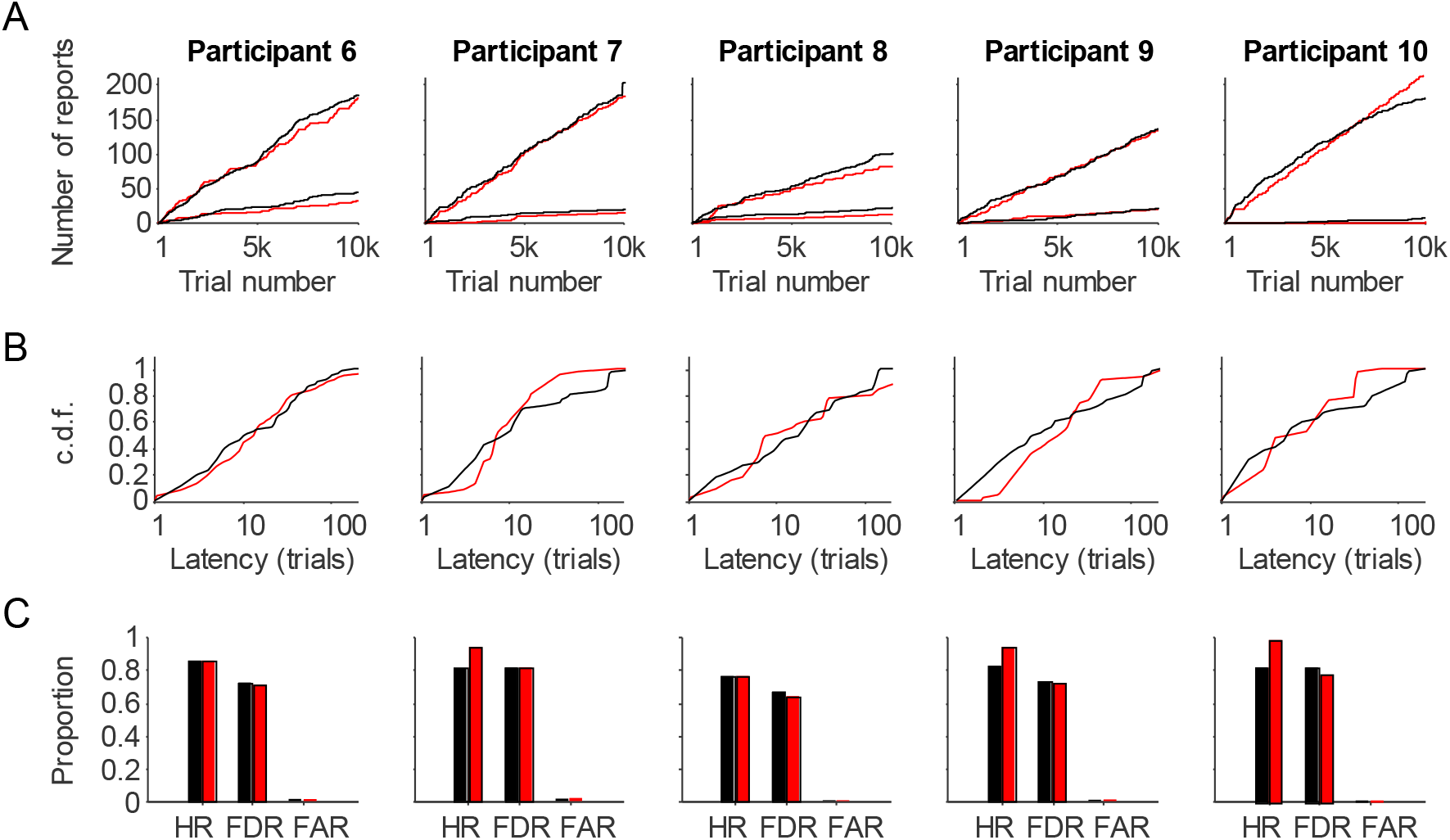
Evaluation of qualitative predictions by the delta-rule model related to detection of changes in the generative function (delta-rule model). The model simulations results (red) were obtained by minimizing the root mean squared deviation (RMSD) with the data (black). (A) Total number of “I think the box has changed” reports (solid) and “I take that back reports (dashed). (B) Cumulative distribution of the number of trials between a change in *p*_true_ and the next “I think the box has changed” report. (C) Hit rates, false discovery rates, and false alarm rates on change point detections. Following Gallistel et al. (2014), the hit rate was defined as the proportion of box changes followed by at least one change report before the next change took place and the false-alarm rate was defined as the number of ‘extra’ change reports (i.e., the number of reports after the first correct report after a change and before the next change) divided by the number of trials on which a change call would have been scored as a false alarm. The false-discovery rate was computed as the proportion of change reports that were ‘extra’ rather than ‘correct’. Results are shown for the (only) five participants in Gallistel et al. (2014) who were asked to report both box changes and second thoughts.

In conclusion, the predictions of a delta-rule model combined with a standard evidence accumulation mechanism are qualitatively consistent with human tracking and detection of changes in the parameter underlying a Bernoulli process. This means that the main argument that Gallistel et al. (2014) presented against delta-rule models does not hold and may stem from an inexhaustive exploration of parameter space.

### Likelihood-based Model Comparisons

The results so far show that just like the IIAB model, the delta-rule model is capable of explaining previously established facts about human performance on probability tracking tasks. But which of the two models explains them *better?* Although the above approach of inspecting summary statistics is useful for checking if a model’s predictions are qualitatively consistent with well-established facts, it cannot be used for quantitative model comparison. The main problem – as also noted by Gallistel et al. (2014) – is that there is no obvious way to weight misestimates in one summary statistic against misestimates in another, which makes it impossible to formulate a single measure to base judgements on.

To compare the models quantitatively, we instead use maximum-likelihood fitting with five-fold cross-validation (see Supplemental Materials for details). We fit the models to the raw data from four experiments (Table 4) reported in three previous studies^4^. The number of trials per participant varied from 9,000 to 10,000 divided over 9 or 10 sessions, with a grand total of 286,890 trials performed by 29 participants over 287 sessions. In each cross-validation fold, a block of 20% of consecutive trials will be left out, which for most datasets amounted to exactly two entire experimental sessions. For now, we limit these analyses to the slider update data because “I think the box has changed” and “I take that back” responses were collected for only some of the participants (we come back to those data below).

**Table 4.** Overview of Datasets Used to Evaluate the Models.

| Exp. ID | Study | Underlying function | Number of participants | Number of trials per participant | Number of trials per session | Total number of sessions |
| --- | --- | --- | --- | --- | --- | --- |
| E1 | Gallistel et al. (2014) | Stepwise | 10 | 10,000 | 1,000 | 100 |
| E2 | Ricci & Gallistel (2017) | Continuous (aperiodic) | 5 | 10,000 | 1,000 | 50 |
| E3 | Ricci & Gallistel (2017) | Continuous (periodic) | 3<sup>5</sup> | 9,000 | 1,000 | 27 |
| E4 | Khaw et al. (2017) | Stepwise | 11 | 9,990 | 999 | 110 |

We first compare the two models contrasted in Gallistel et al. (2014): a single-kernel delta-rule model with a variable response threshold and the IIAB model. We fit the models to all sessions jointly, that is, with a single set of parameters per participant. The delta-rule model accounts for the data better than the IIAB model (Figure 5A): for each of the 29 participants, the delta-rule model is favoured over the IIAB model by a difference of at least 18020 log likelihood points (M±SE: 28654 ± 904)^6^.

**Figure 5.**
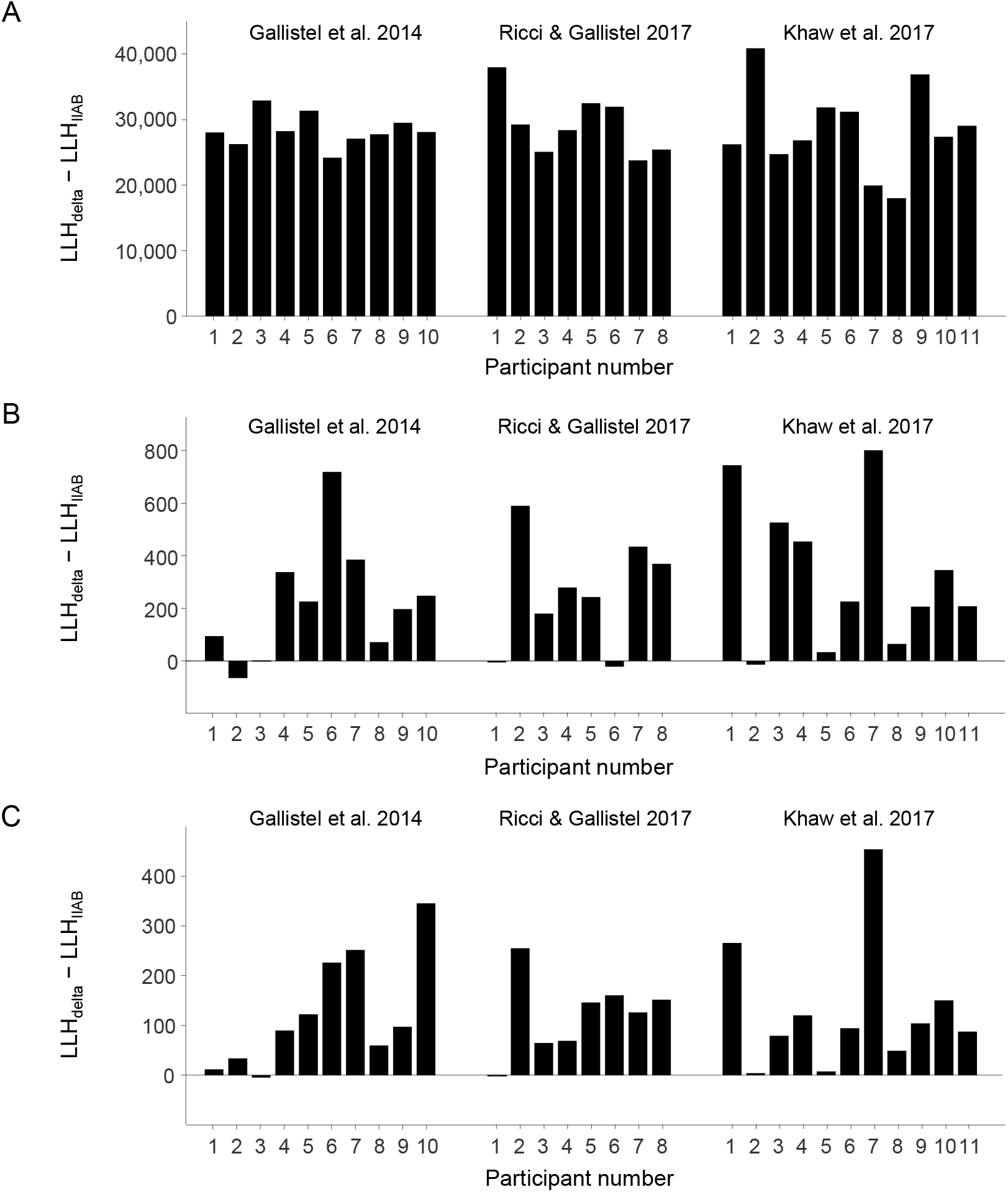
Model comparison results. Model performance is expressed as the log likelihood of the delta-rule model (LLH_delta_) relative to that of the IIAB model (LLH_IIAB_). These values were calculated through five-fold cross validation, by summing the log likelihoods of the left-out trials across the five folds. Positive numbers reflect evidence for the delta-rule model. (A) A delta-rule model with a constrained Gaussian response threshold versus the original IIAB model (without a response threshold). (B) A delta-rule model with a constrained Gaussian response threshold versus an IIAB model with the same response threshold. (C) A delta-rule model with a beta-distributed response threshold versus an IIAB model with the same response threshold.

Hence, not only is the delta-rule model viable from a qualitative perspective, its quantitative account of the raw data is better than that of the alternative model proposed by Gallistel et al. (2014).

There are two big differences between the models that could potentially explain why the IIAB model performs so much worse than the delta-rule model in this comparison: it uses a different mechanism for belief updating and it does not have a threshold on slider reports. To examine how important the assumption of a response threshold is, we next fit a variant of the IIAB model with the same threshold mechanism as in the delta-rule model. This variant has a much better goodness of fit than the original IIAB model (an increase of 370±114 log likelihood points), but it is still outperformed by the delta-rule model for 25 out of 29 participants, with an average log likelihood difference of 271 ± 44 across all participants (Figure 5B). This substantial change in the log likelihood difference suggests that a response threshold is of primary importance to quantitatively account for the data.

A response threshold can be implemented in many ways and which version is chosen can strongly affect the model fit (see Khaw et al., 2017). So far, we have followed Gallistel et al. (2014) by assuming a variable threshold in the shape of a Gaussian distribution with a constraint on the magnitude of the noise. We will now test an alternative version by making two changes. First, we remove the constraint on the amount of variance (*cv*_τ_ ≤ 0.33) because its justification is unclear to us and it may limit both models’ ability to account for participants’ response behaviours. Second, we switch to a beta distribution because, unlike the Gaussian distribution, it produces responses that are bounded between 0 and 1, just like the response scale used by the participants. This change increases the goodness of fit substantially for both the IIAB and delta-rule model, by 650 ± 130 and 495 ± 121 log likelihood points, respectively. The delta-rule model still outperforms the IIAB model for 27 out of 29 participants, with an average difference of 125 ± 20 (Figure 5C). Because the beta distribution provides a better fit, we will employ it in the remaining analyses, unless mentioned otherwise.

So far, we have evaluated the models only based on how well they predict participants’ slider responses. Besides setting the slider, the ten participants in Gallistel et al. (2014) also reported when they thought the box had changed and five of them also reported their second thoughts (using the “I think the box has changed” and “I take that back!” buttons, see Figure 1A). We now also evaluate how well the two models perform when fitting them jointly to slider data and those additional responses (see Supplemental Materials for details and results). We find that both models account reasonably well for the jointly fitted data, with an advantage for the delta-rule-and-evidence-accumulation model in terms of cross-validated log likelihoods: it is the selected model for nine of the ten participants, with a mean difference of 260 ± 74 points. While this shows that bounded evidence accumulation is a viable candidate for explaining change reports in probability estimation tasks, it should be kept in mind that the currently available data is extremely limited (1316 change reports by ten participants and 115 second-thought reports by five participants). Therefore, we view these results mainly as a proof-of-concept and refrain from drawing strong conclusions from them at this point.

In summary, from a quantitative model comparison perspective the delta-rule accounts better for the data than the IIAB model. We checked that this conclusion is robust to changes in the assumed lapse rate, the presence of response noise, the shape of the response noise distribution, and the choice of model comparison method (see Supplemental Materials) and found that the results are highly similar in every test we performed.

### Evaluation of Qualitative Phenomena under Maximum-likelihood Parameters

Likelihood-based model comparison is a powerful tool to evaluate models against each other in a quantitative and principled manner. However, results of such relative comparisons are of little value if none of the models provides a decent account of the data. To verify that this is not the case, we next examine the models’ qualitative predictions under maximum-likelihood parameters. Using these parameter settings, the delta-rule model reproduces the qualitative phenomena related to slider settings almost as well as in the earlier RMSD-based fits (Figure 6). Moreover, it also accounts well for the raw, trial-by-trial slider settings (Figure 7). The maximum-likelihood fits of the original IIAB model (i.e., without response threshold) are very poor (Figures S2 and S3 in Supplemental Materials). After adding a response threshold, the fits become visually of similar quality to those of the delta-rule model (Figures S4 and S5 in Supplemental Materials), which once again highlights that the assumption of a response threshold seems important to account for the data.

**Figure 6.**
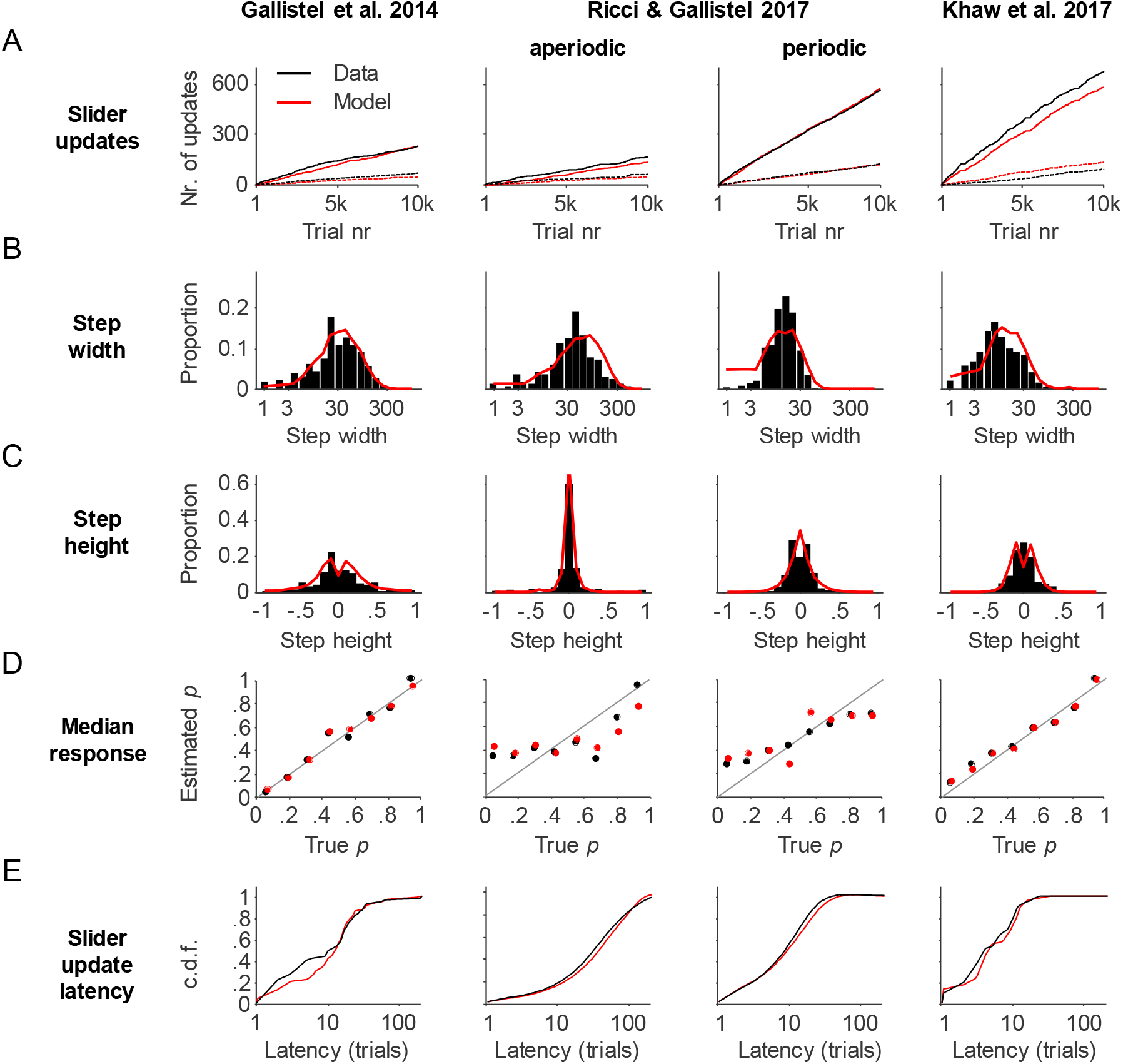
Delta-rule model behavior under maximum-likelihood parameter estimates. Data (black) are shown for Participant 1 in each of the 4 analyzed datasets. The model predictions (red) were obtained by simulating responses using the maximum-likelihood estimates of the paramater values. (A) Total number of slider updates (solid) and number of inconsistent slider updates (dashed) as a function of trial number. (B) Distribution of the number of trials between consecutive slider updates. (C) Distribution of the magnitude of slider updates on trials with an update. (D) Median estimate of the tracked probability versus the median true value. (E) Cumulative distribution of the number of trials between a change in *p*_true_ and the next slider update.

**Figure 7.**
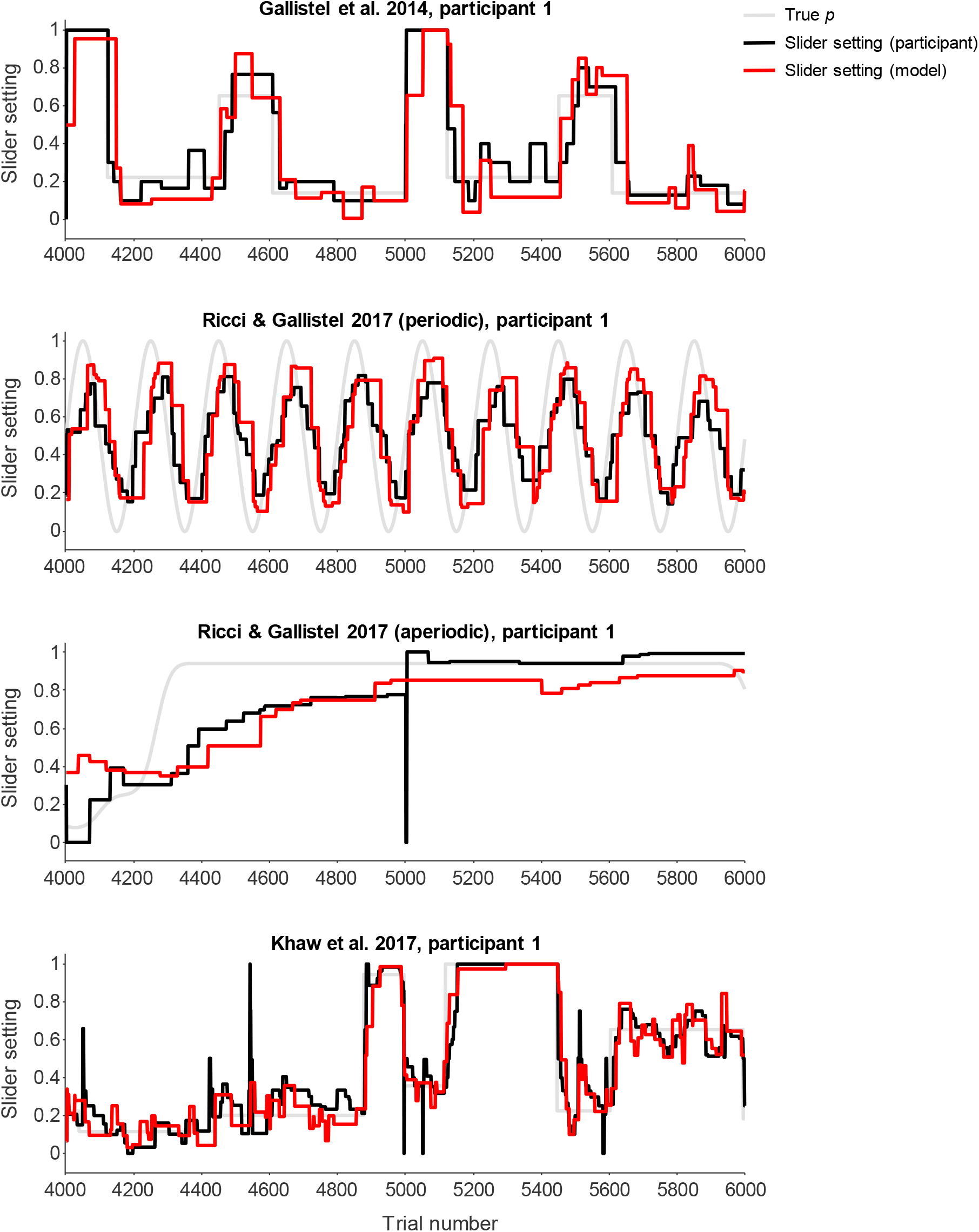
Examples of trial-by-trial slider settings of delta-rule model under maximumlikelihood parameter estimates. For visualisation purposes, only the central 2,000 trials are shown for each dataset.

### Parameter Estimates

#### Response threshold distributions in the delta-rule model

Inspection of the maximum-likelihood estimates of the response thresholds suggests that there is large variation in the trial-to-trial thresholds (Figure 8). As a result, the choice of whether or not to update the slider on any given trial is only partially determined by the discrepancy between the internal belief and the current slider value. Previous literature (Biele, Erev, & Ert, 2009; Gonzalez & Dutt, 2011) has suggested a discrepancy-independent mechanism called “inertia” where the decision to update is entirely determined by the flip of a weighted coin. We tested this mechanism by replacing the response threshold with a constant probability of updating on each trial, implemented as a free parameter. This mechanism makes the fits substantially worse for 27 of the 29 participants, with an average of 69 ± 14 log likelihood points over all participants. This suggests that the update decision at least in part depends on the discrepancy between the internal belief and the current slider value.

**Figure 8.**
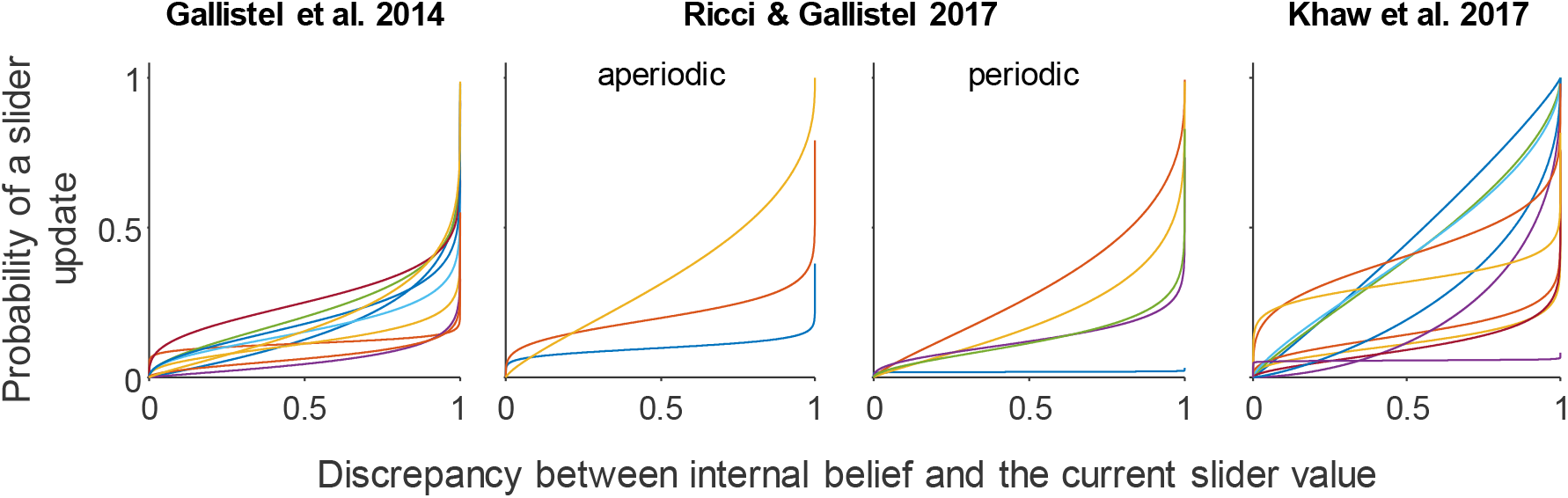
Maximum-likelihood estimates of the variable response thresholds in the delta-rule model (different colors indicate different participants). The threshold is visualised as the cumulative probability distribution of making a slider update as a function of the size of the discrepancy between the internally held belief about the tracked probability and the current slider value. For most participants, the probability of performing a slider update increases with this discrepancy.

#### Response noise in the delta-rule model

The median estimate of the (relative) variance of the beta response noise distribution is 0.058 (IQR: 0.041). To get an intuition of the magnitude of this noise, we performed two simulations. In the first simulation, we generated predicted responses from the delta-rule model for each participant by using the maximum-likelihood parameter estimates. We found that the participant-averaged RMSD between the model predictions and the true value of the tracked probability was 0.192 ± 0.012, which is close to the RMSD of 0.189 ± 0.009 between participants’ slider settings and the true value of the tracked probability. Next, we ‘turned off’ the response noise in the model while keeping all other parameters at the same values. As expected, the RMSD in this simulation was lower (0.165 ± 0.009) but not by much. Hence, the response noise seems to play only a modest role in accounting for participants’ estimation errors.

#### Decision threshold in the IIAB model

The decision threshold parameter in the IIAB model – which controls when the model considers the current belief to be “broke” and in need of an update – is estimated to be close to 0 for every participant (*M*=0.032, *SE*=0.018). Importantly, this means that the IIAB model captures the data best when setting its parameters in such a way that it considers its current probability estimate “broke” on each trial and thus transforms into a trial-by-trial updating model in which the stepwise behaviour is entirely explained by the response threshold.

### Slider Updating Consistency

Why do people regularly make a slider update that is inconsistent with their last observation, such as decreasing their estimate of the probability of red outcomes after observing a red outcome? In a basic delta-rule model, response updates are always consistent with the most recent observation: observing a red ring increases the estimate of the probability of observing a red ring and observing a ring of the other colour decreases it. In the IIAB model, the “second thoughts” mechanism might on rare occasions cause inconsistent updating (4.50 ± 0.65% of all slider updates under the maximum likelihood parameter values).

One potentially important source of inconsistent updating is the response threshold. For example, a momentarily high threshold might suppress a downwards adjustment of the slider but it will never suppress a downwards adjustment of the internal belief. If the threshold on the next trial happens to be lower, and the new observation increases the internal belief by less than it was decreased on the previous trial, the reported estimate will be adjusted downwards – which would be inconsistent with the last observation. Indeed, the maximum-likelihood fits of the IIAB model and delta-rule model with a response threshold predict that 31.8±0.8% and 22.1±0.9% of the slider updates, respectively, are inconsistent. Hence, the IIAB model slightly overestimates the empirical proportion of 23.6±1.6% (BF_10_ = 305; two-tailed paired-samples *t*-test), while the predictions of the delta-rule model are consistent with the data (BF_10_ = 0.36).

## DISCUSSION

Previous studies where participants track a non-stationary Bernoulli distribution (Gallistel et al., 2014; Khaw et al., 2017; Ricci & Gallistel, 2017; Robinson, 1964) have consistently observed stepwise, “staircase-like” response patterns. It has been claimed that this pattern and related phenomena are inconsistent with models from the associationist tradition and are instead indicative of discrete, stepwise learning through hypothesis testing (Gallistel et al., 2014; Ricci & Gallistel, 2017). If correct, this claim would constitute a serious challenge to the neuropsychological literature which connects trial-by-trial learning of probabilities (Nassar et al., 2012, 2010; Norton et al., 2019; Wilson et al., 2013, 2018), encoding of prediction errors in the anterior cingulate cortex (Behrens et al., 2007; Rushworth & Behrens, 2008; Silvetti et al., 2013) and the experience of surprise (Lavín, Martín, & Jubal, 2014; Preuschoff, ’t Hart, & Einhäuser, 2011).

In the present paper, we argue that the rejection of associative learning in human probability estimation was premature because it was based on an incomplete investigation of the predictions made by delta-rule models: parameter space was explored manually and no model fitting was performed. To reassess the earlier drawn conclusions, we reanalysed data from three previous studies (Gallistel et al., 2014; Khaw et al., 2017; Ricci & Gallistel, 2017) using rigorous model fitting and model comparison methods. Our findings demonstrate that a dual process of two broadly supported computational theories – the delta-rule for online learning of a latent variable and bounded evidence accumulation for making categorical decisions – makes predictions that are qualitatively highly consistent with the observed phenomena. Moreover, quantitative model comparison showed that this combination of established theories actually accounts *better* for the data than the IIAB hypothesis-testing model that was specifically developed for this task. These conclusions hold across all tested data sets and are robust to changes in the modelling assumptions about the shape of the response threshold distribution, the assumed lapse rate, the presence of response noise, and the choice of model comparison method. We thus conclude that associative models of probability estimation are still viable candidates. However, we do not take this to imply that there is no place for hypothesis testing models. On the contrary, our point is that the experimental task requires both *descriptive tracking* of the observed frequencies and *inferences to unobservable states* of the world (hypotheses about whether the box has changed). The mind can do both and previous research has suggested models of those processes. Our central claim here is that these previous models can explain participants’ ability to “do both” in the present task too.

We note that we are not the first to combine associative learning and evidence accumulation decision making.^7^ Fontanesi et al. (2019) have shown how drift-rates can be updated associatively in a multi-armed bandit task. Like them, we believe that this kind of approach might prove fruitful: take processes that we know the mind is capable of and investigate how they together might generate more complex behaviour.

### Theoretical importance and implications of a variable response threshold

Adding a variable response threshold greatly improved the fits of both models. One reason is that participants make inconsistent updates which are incompatible with the original models. The variable response threshold allows the models to account for this. Hence, this threshold does not merely “soak up noise” but is required by both the IIAB and delta-rule model to explain inconsistent updating and other empirical phenomena (Table 1). We therefore emphasise that a variable response threshold does not represent a “nuisance term”, akin to adding an error term to a regression, but constitutes a theoretical proposition which is tentatively supported by our results.

Evaluation of the fitted response thresholds revealed that many distributions were so broad that the choice of whether or not to update on any given trial becomes partly stimulus-independent. Completely stimulus-independent thresholds have elsewhere (Biele et al., 2009; Gonzalez & Dutt, 2011) been termed “inertia”. For two of the 29 participants, a coin-flip mechanism did indeed provide a better quantitative fit than the response threshold mechanism. However, for the vast majority of participants it did not, which suggests that updating is at least in part driven by stimulus-dependent factors (as also concluded by Khaw et al., 2017). For other participants, we obtained threshold distributions such that the probability of updating the response increased with the discrepancy between the current response and the internal estimate. Updates were less likely under very small discrepancies and more likely under very large discrepancies. We interpret this as a *resistance* to updating, as opposed to a suppressive *threshold* – the term we have hitherto used. Participants are reluctant to update – perhaps due to the motor or time cost – but balance this against their wish to respond correctly. They care about not being *very* wrong, but not so much about being *exactly* right. In economics, the idea that learning can be influenced by a trade-off between the costs of updating and the gains from a more accurate belief has been formalised in the “rational inattention” literature (Sims, 2003). The stepwise response pattern in the present Bernoulli distribution task has been taken to support this idea (Khaw et al., 2017). The reluctance interpretation of our threshold distributions stated above is different to rational inattention in that it supposes that the overt *response*, and not the covert *estimate*, is affected by the trade-off. Our modelling here was not aimed at answering which version is correct, which could be an interesting avenue for future work.

Inertia, resistance, and rational inattention are distinct theoretical propositions as to how the mind times response updates. It may be that there is true heterogeneity in what mechanisms are used or that there exists a single mechanism which can express itself in (ostensibly) different ways. Regardless of how internal estimates are updated, the process which mediates their expression as overt behaviour is scientifically interesting in itself and deserves further attention.

### Observation weighting is intrinsic to the theories

We equalised the delta-rule model and the IIAB model on the assumption of a variable response threshold to show that this, although important, is not what drives the conclusions. Another difference is that the delta-rule model effectively performs *unequal weighting* of *all* observations while the IIAB model performs *equal weighting* of an *ordered subset* of observations (those that occurred since the last or second to last change point). The weighting schemes are defining features of the theories the models embody. The IIAB model implements the theory that “the perception of Bernoulli probability is a by-product of the real-time construction of a compact encoding of the evolving sequence by means of change points” (Gallistel et al., 2014). Under unequal weighting of observations, the model would contradict this theory – the percept would no longer be deduced from the change points. Associative theories instead suppose that the percept is not a by-product, but *learned in itself by* the gradual adjustment of an associative value. The way that observations are weighted thus cannot be equalised across models without fundamentally changing the theories that one is contrasting.

### Alternative models

The present work focused on the question of whether Gallistel et al., (2014) were right in their rejection of associative learning in human perception of non-stationary probabilities. We did so by focusing on the standard delta-rule model and comparing it with the alternative proposed by Gallistel et al., (2014). However, there is a myriad of possible variations on these models that may also be worth to consider when addressing the more general question of how to best explain the data.

One such candidate, which was also tested by Gallistel et al., (2014) with manually selected parameter values, is a two-kernel version of the delta-rule model. This type of model continuously entertains two estimates (typically one based on a slow learning rate and another one based on a fast learning rate) and selects one to report on each trial. Preliminary results (see Supplemental Materials) suggest that this kind of model performs better than the single-kernel version. A related variant, that has been proposed in the literature but which we did not test here, is a delta-rule model with a learning rate that adapts as a function of the prediction error modulated by the estimated volatility (Behrens et al., 2007; McGuire, Nassar, Gold, & Kable, 2014) or declarative information about the choice environment (Lee, Gold, & Kable, 2020). Future studies might want to pool a larger number of datasets (from non-Bernoulli distribution tasks too) and compare various adaptive learning rate models to multi-kernel models. One interesting prediction from multi-kernel models is that people should be able to report several earnest estimates at any one point, which should be possible to observe in an experiment.

Another interesting variant to consider is the ‘approximately Bayesian’ delta-rule model from the neuropsychological literature (Nassar et al., 2010). Preliminary results (see Supplementary Materials) suggest that this model performs better than the hypothesis testing model but worse than the regular delta-rule. Allowing it to underweight likelihoods helps, in line with a previous observation (Nassar et al., 2010). However, with this change the model’s original theoretical claim (that people are approximate Bayesians who balance two hypotheses trial-by-trial) becomes less distinct from the more general notion of the learning rate being inconstant. Our tentative interpretation is that the common problem of the (normative) Bayesian delta-rule and the IIAB is not that they adapt observation weights (which is supported by other evidence, see Behrens et al., 2007; Krugel et al., 2009) but could be that they do this by considering a limited number of discrete hypotheses.

Finally, Costello and Watts (2014, 2016, 2018) have suggested that a range of results from various probability judgement and decision tasks, including the present paradigm, could arise from normatively correct judgements being perturbed by constant memory noise. They simulated a hypothesis testing model (Costello & Watts, 2018) with the same two stages/three hypotheses structure as the IIAB. They argue that, if there is constant memory noise, updates from re-estimation will be biased towards 0.5 and updates from acceptance of a new hypothesis will be biased towards the extremes. These effects should cancel out, making the estimates accurate on average (phenomenon 4, Table 1). If descriptive estimates are actually made associatively, and hypothesis testing is a separate evidence accumulation process, Costello and Watts’s (2018) framework predicts a constant bias towards 0.5, which seems inconsistent with the available data.

### Interference from distal beliefs as an alternative to proximal learning rate adaptation

Previous research has suggested various mechanisms for adapting the learning rate as a part of the associative process. Our combined model highlights a complementary solution. One can imagine a process where the proximal learning not only feeds evidence “upwards” to the accumulator that affects beliefs, but where beliefs reached by the accumulator are also signalled back “down” to the online learning which adapts to new beliefs about the world. For example, when a decision has been made that the box has changed it could prompt a momentary increase in the learning rate to adapt to the new reality. Lee et al. (2020) found that people adapted their learning rate depending on verbal information about the generative function. Using our vocabulary, proximal learning in their task was affected by distal beliefs informed by written instructions. Our conjecture here is that a similar effect might be observed when the distal beliefs (what Lee called the “cognitive map”) are arrived at through experience.

In the approximately Bayesian delta-rule model (Nassar et al., 2010) the perceived probability that there has been a change point determines the learning rate. In other words, it explains learning rate adaptation as part of the associative process. By contrast, we speculate that delta-rule learning might be interfered with by semantic cognition (Rogers & McClelland, 2008). These ideas are not mutually exclusive, but the latter seems necessary to satisfactorily account for any effects of external information. Outside the laboratory, we often have a rich semantic understanding of the generative processes we “draw” from. Future work could consider how such information can affect online estimation.

### Unexplained phenomena

Phenomena 10 and 11 (Table 1) cannot be explained by either the IIAB or the delta-rule model as implemented here. However, both models could in principle be extended to do so.

Gallistel et al. (2014) noted that participants’ frequency of change point reports on average decreased per session (phenomenon 10). They concluded that the IIAB cannot explain this under the regular priors used to explain the other qualitative phenomena, and that they had to substitute special priors tailored to this summary statistic. It is, however, easy to imagine a process where the threshold in the troubleshooting stage, *T*_2_, is not fixed but adapted over time. This would result in a changing change point detection frequency. Analogously, the evidence accumulation literature explains this kind of effect as decision bound separation being adapted through learning (Liu & Watanabe, 2012; Zhang & Rowe, 2014).

In Ricci and Gallistel (2017), some participants were able to correctly report having drawn from a sinusoidal during the debriefing (phenomenon 11). A central theoretical proposition of the IIAB (see pp. 106, Gallistel et al., 2014) is that people do not perceive probabilities per se but “deduce” them from a (sparse) memory of change points. To generate a declarative belief of a continuous functional form from a discrete set of memories, the IIAB would require some function learning mechanism (e.g., Brehmer, 1974) which interpolates between the “datapoints”. For the delta-rule model, we need the mechanism to be recursive. There exist several recursive function learning models, some of which are specifically adapted to non-stationary environments (Speekenbrink & Shanks, 2010) and some of which use a version of delta-rule learning (DeLosh, Busemeyer, & McDaniel, 1997). The perhaps most famous of the latter is the EXAM model (Mcdaniel & Busemeyer, 2005).

In sum, we do not view phenomena 10 and 11 as evidence against either model but rather as avenues of future research. Investigating phenomenon 10 involves opening a black box by trying to establish a structured explanation of aspects which we here model as free parameters. Investigating phenomenon 11 would involve attaching a third process of function learning to what we suggest could be a dual process of delta-rule online learning and evidence accumulation decision making.

### Neuropsychological predictions

Since both associative learning and evidence accumulation are established processes, there already exists a neuropsychological literature which can help us (tentatively) device experimental tests of our combined model. Evidence accumulation might be mediated in the anterior dorsal striatum (ADS: Yartsev et al., 2018) while the eventual discrete decision making seems to be mediated by the prefrontal cortex (Philiastides, Auksztulewicz, Heekeren, & Blankenburg, 2011). The evidence our model accumulates is positive and negative (directional) prediction errors, which it also uses to update the online estimate. Such prediction errors are encoded separately in the anterior cingulate cortex (ACC: Rushworth & Behrens, 2008) while dopamine neurons in the ventral striatum (O’Doherty et al., 2004) and midbrain (Bayer & Glimcher, 2005) only seem to encode positive prediction errors while negative prediction errors yield an absence of activity. If the present task is replicated while activity in these regions of interest (ROI) is monitored, it should be possible to connect activity in each ROI to the applicable behavioural statistic (as done by Behrens et al., 2007, for the ACC in a related task).

### Concluding remarks

Previous research has highlighted that a complete theory of probability perception must account for both online estimates of the proximal frequencies and detection of changes in the distal generative process. We have demonstrated that it was premature for the previous literature to rule out a role for associative models in this. We have shown here that the perception of probability is consistent with a combination of mechanisms from both the empiricist and rationalist traditions that are already well-established in the literature.

## Footnotes

1 We are grateful to two anonymous reviewers for helping us think more clearly about this.

2 In addition to a belief (hypothesis) about the underlying probability, the model also assumes a belief about the probability that the underlying probability is changed on the next trial. Because that “change probability” has no important role in our discussion of the model, we will not discuss it further here.

3 For example, in the experiment by Gallistel et al. (2014), trials with an update took on average three times longer (4.22 ± 0.18 seconds) than trials without an update (1.39 ± 0.01 seconds). Responding on each trial would almost have tripled the median session time – from around 25 minutes to around 70 minutes.

4 There is one other study using the same paradigm (Robinson, 1964), but it has no preserved record of the data known to us.

5 This experiment had 4 participants, but we suspect that for one of them the responses were flipped between two sessions. We excluded this participant from our analyses.

6 When fitting the models separately to each session, the average difference is 285±40 in favour of the delta model. Considering that log likelihoods scale linearly with the number of trials, this difference is comparable to that obtained by fitting the full datasets.

7 We are grateful to Mona Guath for making us aware of this.

## Notes

We have no conflicts of interest to disclose. The research was funded by the Swedish Research Council (Grant 2018-01947) and the Marcus and Amalia Wallenberg Foundation (MAW 2016.0132). Earlier versions of this article were disseminated on Biorxiv as preprints.

https://osf.io/zhv2r/

### Competing Interest Statement

The authors have declared no competing interest.

### Summary of Updates

We further clarified the conceptual background and the logic of our paper in multiple ways; We explain more clearly how we deal with the issue of model flexibility (cross validation); We now also fit the change reports and second thought reports using maximum likelihood fitting.

