## Supplemental Text and Figures for "Further Perceptions of Probability: In Defence of Associative Models – A Commentary on Gallistel et al. 2014"

### SUPPLEMENTAL MATERIALS

#### Likelihood function

The likelihood function for models of psychophysical tasks generally takes the form

$$L(\boldsymbol{\theta}) = p(\mathbf{R} \mid \mathbf{S}, \boldsymbol{\theta}),$$

where $\mathbf{R}=\{R_1, R_2, \ldots, R_N\}$ is a vector with responses for all trials 1 to $N$, $\mathbf{S}=\{S_1, S_2, \ldots, S_N\}$ is a vector with the presented stimulus values, and $\boldsymbol{\theta}$ is a vector with parameter values. In the present experiment, $R_t$ is the slider value on trial $t$ and $S_t$ is the observed Bernoulli outcome (e.g., “green” or “red”) on trial $t$. Assuming that trial responses are conditionally independent and do not depend on future events, we can rewrite this to

$$L(\boldsymbol{\theta}) = \prod_{t=1}^{N} p(R_t \mid S_{1\ldots t}, \boldsymbol{\theta}).$$

For many psychophysical tasks, it can be safely assumed that the response on trial $t$ depends only on the stimulus presented on that trial – rather than the entire stimulus history – which allows to greatly simplify the likelihood function by factorising it. In the present experiment, we cannot make that assumption, because the response on trial $t$ depends not only on the outcome (“green” or “red”) on that trial, but also on the stimulus outcomes of preceding trials. Fortunately, however, the stimulus sequence becomes irrelevant once we know the model’s current internal estimate of the tracked probability, $\hat{p}_{\mathrm{B},t}$. On trials where the discrepancy between the internal estimate of the tracked variable and the current slider value does not exceed the response threshold, $\Delta_t \equiv |\,\hat{p}_{\mathrm{B},t} - R_{t\text{-}1}| \le \varepsilon$, the model predicts no slider update, such that $p(R_t \mid \Delta_t \le \varepsilon) = \delta(R_t - R_{t\text{-}1})$, where $\delta$ is the Dirac delta function. On trials where $\Delta_t$ *does* exceed the response threshold, the model predicts the slider update to be a draw from a beta distribution (see section *Response noise* in main text), such that $p(R_t \mid \Delta_t > \varepsilon) = f_\beta(R_t; \mu_\beta = \hat{p}_{B,t}, V_B)$, where $f_\beta(x; \mu, V)$ is the probability density function of the beta distribution with a mean $\mu$ and variance $V$, evaluated at $x$.

The likelihood function can now be evaluated in a rather straightforward way as follows. First, use a forward simulation to compute the model’s internal estimates $\hat{p}_{\mathrm{B},1}, \ldots, \hat{p}_{\mathrm{B},N}$ for the entire trial sequence, which are uniquely determined by the stimulus sequence and model parameters. Next, compute for each trial the discrepancy $\Delta_t$ between the internal estimate, $\hat{p}_{\mathrm{B},t}$, and the participant’s slider value at the start of the trial, $R_{t\text{-}1}$. Then, compute for each trial the probability $U_t$ that an update will take place, that is, the probability that $\Delta_t$ exceeds a random draw from the distribution that specifies the variable response threshold. Ignoring lapses, the probability density function for the participant's response on trial *t* now takes the form

$$p_{\text{no lapse}}\left(R_t \mid \Delta_t, U_t\right) = U_t\, p\left(R_t \mid \Delta_t > \varepsilon\right) + \left(1 - U_t\right) p\left(R_t \mid \Delta_t \le \varepsilon\right).$$

In all experiments, we discretized the response scale to $n_{\text{bins}}$=100 values linearly spaced in [0, 1]. Including the lapse rate, *L*, and using the midpoint rule for integration, the probability of the response for trial *t* is computed as

$$p\left(R_t \mid \Delta_t, U_t\right) = \left(1 - L\right) p_{\text{no lapse}}\left(R_t \mid \Delta_t, U_t\right) \frac{1}{n_{\text{bins}}} + L \frac{1}{n_{\text{bins}}}.$$

**Parameter fitting and model comparison methods**

We use the Bayesian Adaptive Direct Search (BADS) algorithm (Acerbi & Ma, 2017) to find for each participant the parameter values that maximise the log of the likelihood function. In order to reduce the risk that the algorithm terminates in a local maximum, we run it thirty times with different initial parameter values. Prior to each run, we evaluate the likelihood function for five hundred randomly drawn parameter vectors and choose the vector with the highest likelihood value as the initial parameter vector for BADS. Finally, to compare models, we use five-fold cross-validation. In the first of the five runs, the first 20% of the trials are left out during parameter fitting; in the second run, the second 20% of the trials are left out; etc. In each run, we compute the log likelihood on the left-out trials based on parameters fitted to the remaining 80% of the trials. We sum the log likelihood values across the five runs to obtain a single "cross-validated log likelihood" value per fit.

**Model recovery**

To verify that our modelling methods can accurately recover parameter values and allow for reliable model comparison, we performed a model recovery analysis. We created eight synthetic data sets from the IIAB and the delta-rule model, both extended with a beta-distributed response threshold. This was done by simulating responses based on the maximum-likelihood parameter values of the first two participants in each of the four datasets. The stimuli in these simulations were the same as those presented to the respective participants. Next, we used the method described above to fit the two models to all sixteen datasets. The results show good model selection performance (Figure S1A), which indicates that the models make clearly distinguishable predictions and that our model comparison methods successfully detect these differences. Moreover, we find that parameter values are recovered accurately (Figure S1B), which again indicates that our methods are adequate. More importantly, this also indicates that the mechanisms behind the parameters have clearly distinguishable effects on the model predictions; if, for example, the variable response threshold would affect the model predictions in the same way as the Stage-1 threshold in the IIAB model, then we would not have been able to accurately recover the generative parameter values for these two mechanisms.

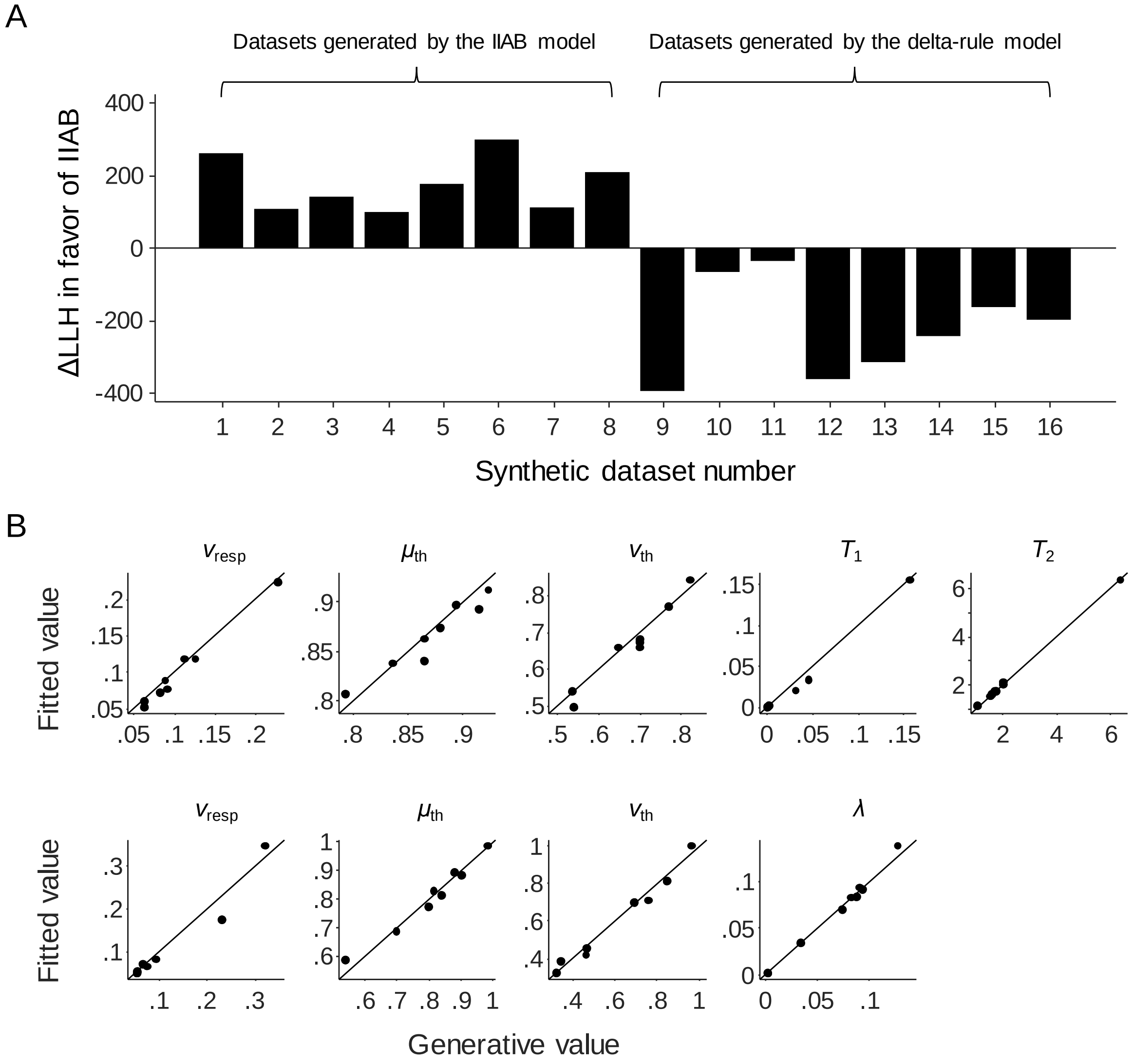


**Figure S1 | Model recovery results.** (A) Model comparison results on 8 synthetic datasets generated by the IIAB model (sets 1-8) and 8 synthetic datasets generated by the delta-rule model (sets 9-16). On all 16 datasets, the cross-validated log likelihood for the generating model was higher than that of the alternative model, which indicates good model selection performance. (B) Estimated parameter values for the synthetic datasets generated by the IIAB model (B) and the delta-rule model (C). For all 72 parameter estimates, the fitted value is close to the generative value. This indicates that the model parameters have largely independent effects on the predictions and that parameter estimation established by our methods is accurate.

**Maximum-likelihood fits of change report data**

Participants in the experiment by Gallistel et al. (2014) were asked to not only estimate the probability underlying the observed rings, but also report when they suspected that the probability had changed ("I think the box has changed") and when they had second thoughts about those reports ("I take that back"; the latter type of response was recorded for only five of their ten subjects). Estimating the probability (slider settings) and detecting changes in the box ("I think the box has changed" and "I take that back" reports) are separate tasks that could be driven by separate mechanisms. The results in the main text showed that a model based on delta-rule learning accounts well for the slider settings. In the present section we test if a standard bounded evidence accumulator model can account for the change reports and second thoughts.

To this end, we extend the delta-rule model with an accumulator on the trial-by-trial prediction error, as described in the Modelling section in the main text. Each time when the accumulated error crosses the decision bound (a free parameter), the model generates a change report ("I think the box has changed"). After each change report, a temporary second decision bound is activated. This bound is at the same position as the change report bound but in the opposite direction to the detected change point (see Figure 2, main text). The lifetime of the second bound is another free parameter. If the second bound is crossed before it expires, a second thought report ("I take that back") is recorded.

One problem with the predicted change reports in both the IIAB and delta-rule model is that they are deterministic: on each trial there is a change report or there isn't, and there either is a second though report or there isn't. This is problematic from a model fitting perspective, because one wrong prediction will reduce the likelihood of each model to 0. Put differently, the data are 'fatal' to both models unless they are 100% accurate in their change point and second thought predictions, which is an unrealistic expectation. To solve this, we turn the deterministic predictions of both models into stochastic ones by adding Gaussian noise to the timing of the predicted responses:

$$
\begin{aligned}
&p\left(\text{change report on trial } x \mid \mathbf{Y}\right) = \sum_{i=1}^{n_{\text{cp}}} N\left(x; Y_i, \sigma\right) \\
&p\left(\text{second thought on trial } x \mid \mathbf{Z}\right) = \sum_{i=1}^{n_{\text{st}}} N\left(x; Z_i, \sigma\right),
\end{aligned}
$$

where $\mathbf{Y} = \left\{ Y_1,\ Y_2,\ ...,\ Y_{n_{\mathrm{cp}}} \right\}$ and $\mathbf{Z} = \left\{ Z_1,\ Z_2,\ ...,\ Z_{n_{\mathrm{st}}} \right\}$ are the trial numbers on which the model predicted change reports and second thoughts, respectively; $\sigma$ is a free parameter that controls the amount of uncertainty in the predicted timings (with $\sigma$=0 being the special case of deterministic responses); $N(x;\ \mu,\ \sigma)$ is the normal distribution with mean $\mu$ and standard deviation $\sigma$, evaluated at $x$.

We used maximum-likelihood estimation with cross validation to fit the parameters related to the change reports to the data from the participants in Gallistel et al. (2014). For computational efficiency, the other model parameters were fixed to the maximum-likelihood values that we found earlier when fitting the models to the slider responses. The results show that the extended delta-rule model captures the empirical trends quite well (Figure S2). Moreover, model comparison based on cross-validated log likelihood values shows that the extended delta-rule model is preferred over the IIAB model for nine of the ten participants, with an average difference of 260 ± 74 points.

**IIAB model fits**

Figures S3 and S4 show cross-validated maximum-likelihood fits of the original IIAB model, that is, without a response threshold[1]. This model provides a poor account of the summary statistics. The quality of the fits increases substantially once we add a response threshold (Figures S5 and S6).

[1] We did, however, include response noise and a lapse rate in our implementation of this model, to avoid numerical instabilities

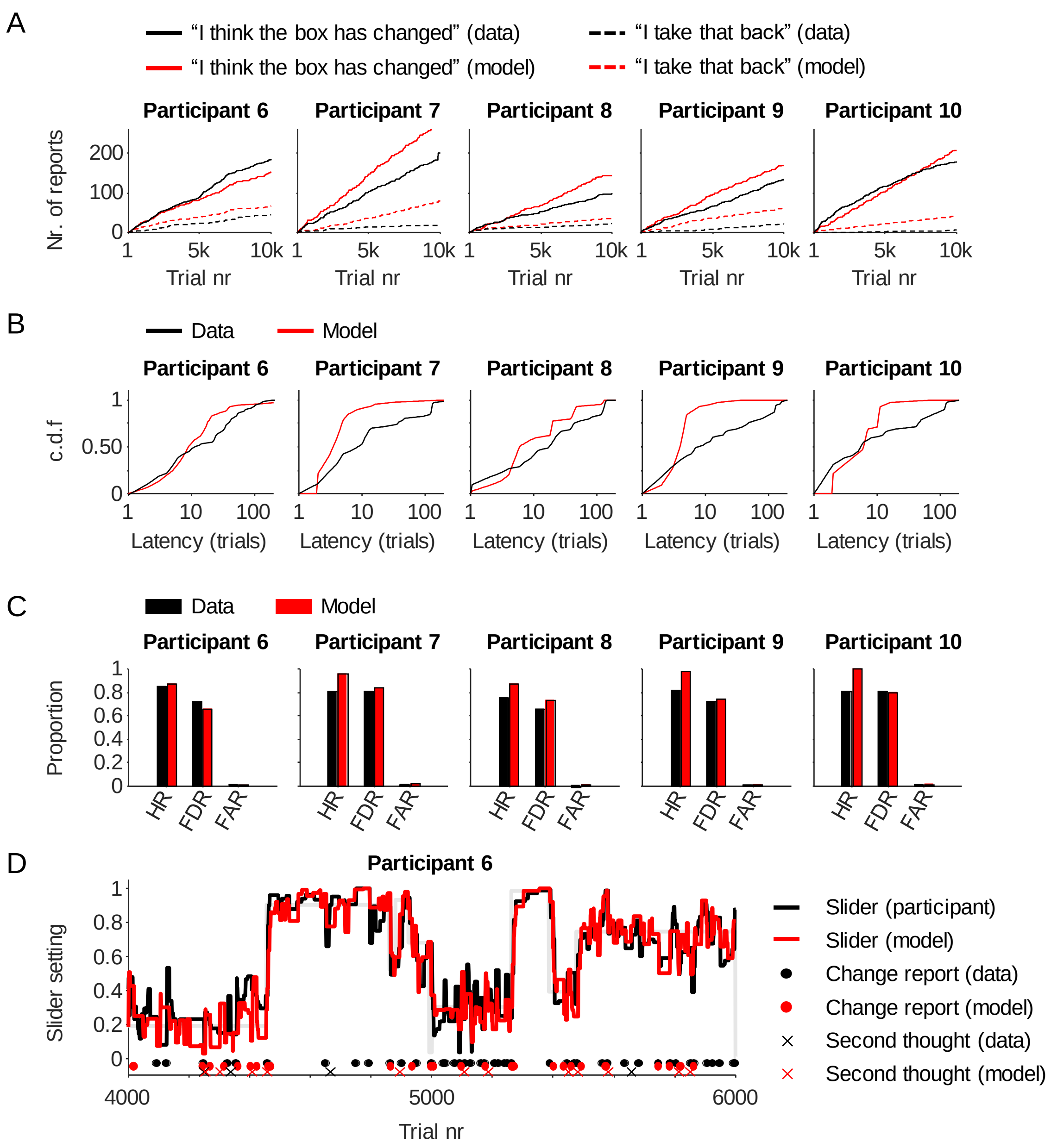


**Figure S2 | Maximum-likelihood fits to phenomena related to change reports and second thought reports.** The plots show summary statistics related to the change reports ("I think the box has changed") and second thoughts ("I take that back") that were recorded for five participants in the experiment by Gallistel et al. (2014). Also shown are fits from the delta-rule model extended with a bounded accumulator mechanism to make predictions about change reports and second thoughts. (A) Cumulative number of change reports and second thougths. (B) Cumulative distribution of change report latencies, that is, the number of trials between a change in the box and the participant's first subsequent change report. (C) Hit rates, false-discovery rates, and false-alarm rates. Following Gallistel et al. (2014), the hit rate was defined as the proportion of box changes followed by at least one change report before the next change took place and the false-alarm rate was defined as the number of 'extra' change reports (i.e., the number of reports after the first correct report after a change and before the next change) divided by the number of trials on which a change call would have been scored as a false alarm. The false-discovery rate was computed as the proportion of change reports that were 'extra' rather than 'correct'. (D) Example of empirical and predicted change reports and second thoughts (central 2000 trials of participant 6).

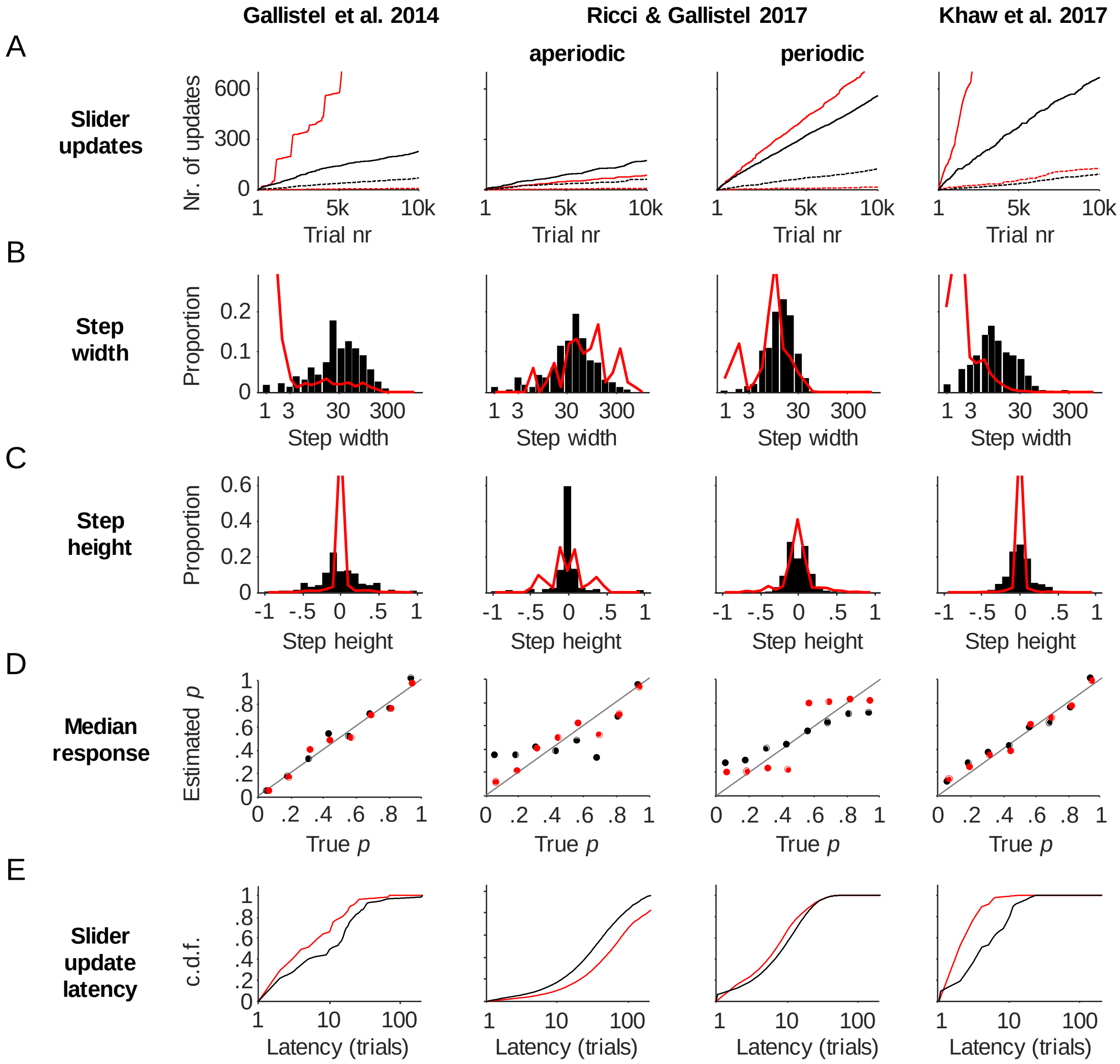


**Figure S3 | Maximum-likelihood fits of the original IIAB model.** Results are shown for Participant 1 in each of the 4 analyzed datasets. The model simulations results were obtained by simulating responses from the model using the maximum-likelihood estimates of the parmater values. (A) Total number of slider updates (solid) and number of inconsistent slider updates (dashed) as a function of trial number. (B) Distribution of the number of trials between consecutive slider updates. (C) Distribution of the magnitude of slider updates on trials with an update. (D) Median estimate of the tracked probability versus the median true value. (E) Cumulative distribution of the number of trials between a change in $p_{\text{true}}$ and the next slider update.

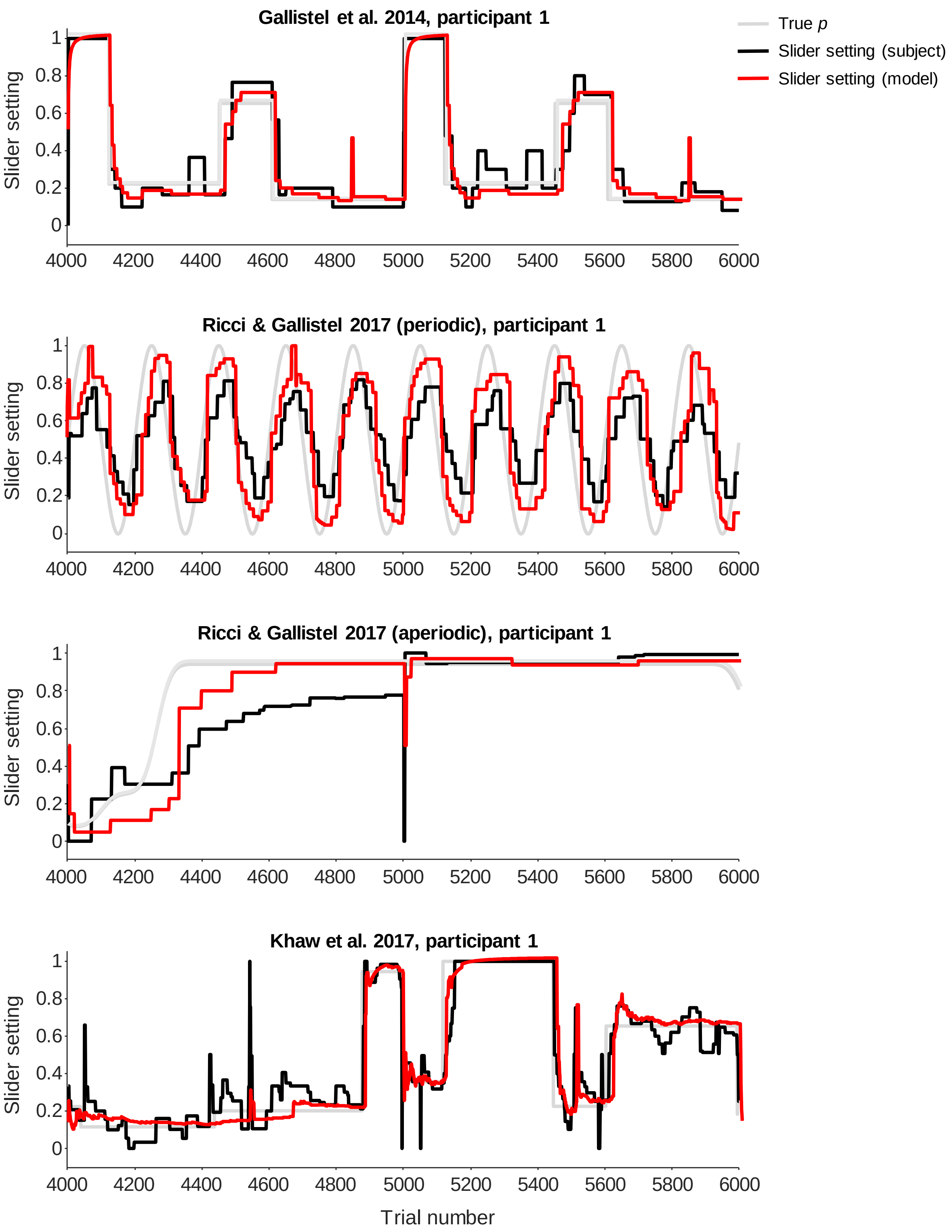


**Figure S4 | Examples of trial-by-trial slider settings of the original IIAB model.** The predictions were generated by simulating responses from the model under maximum-likelihood parameter estimates. For visualisation purposes, only the central 2,000 trials are shown for each dataset.

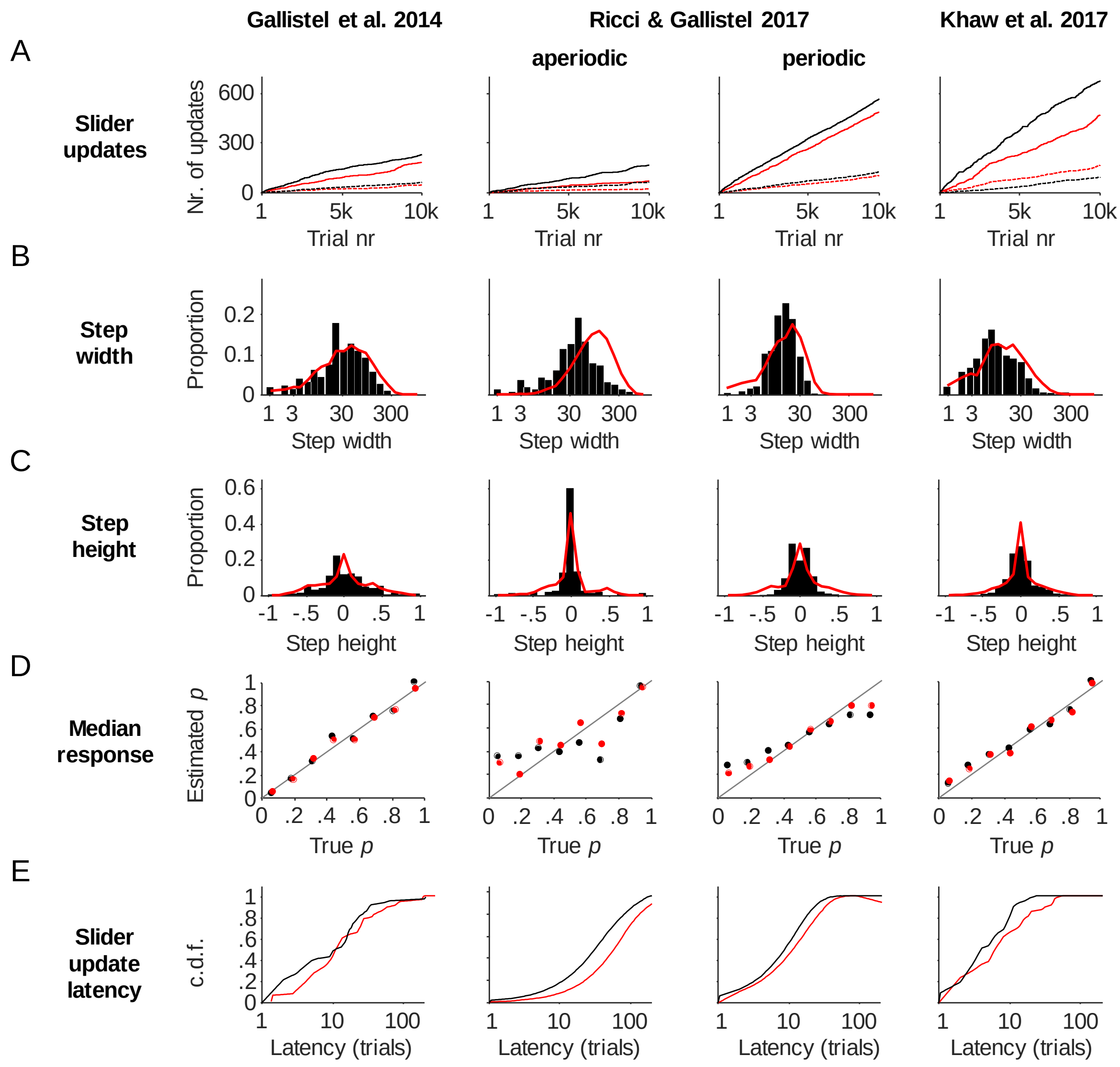


**Figure S5 | Maximum-likelihood fits of the IIAB model extended with a variable response threshold.** Results are shown for Participant 1 in each of the 4 analyzed datasets. The model simulations results were obtained by simulating responses from the model using the maximum-likelihood estimates of the parmater values. (A) Total number of slider updates (solid) and number of inconsistent slider updates (dashed) as a function of trial number. (B) Distribution of the number of trials between consecutive slider updates. (C) Distribution of the magnitude of slider updates on trials with an update. (D) Median estimate of the tracked probability versus the median true value. (E) Cumulative distribution of the number of trials between a change in $p_{\mathrm{true}}$ and the next slider update.

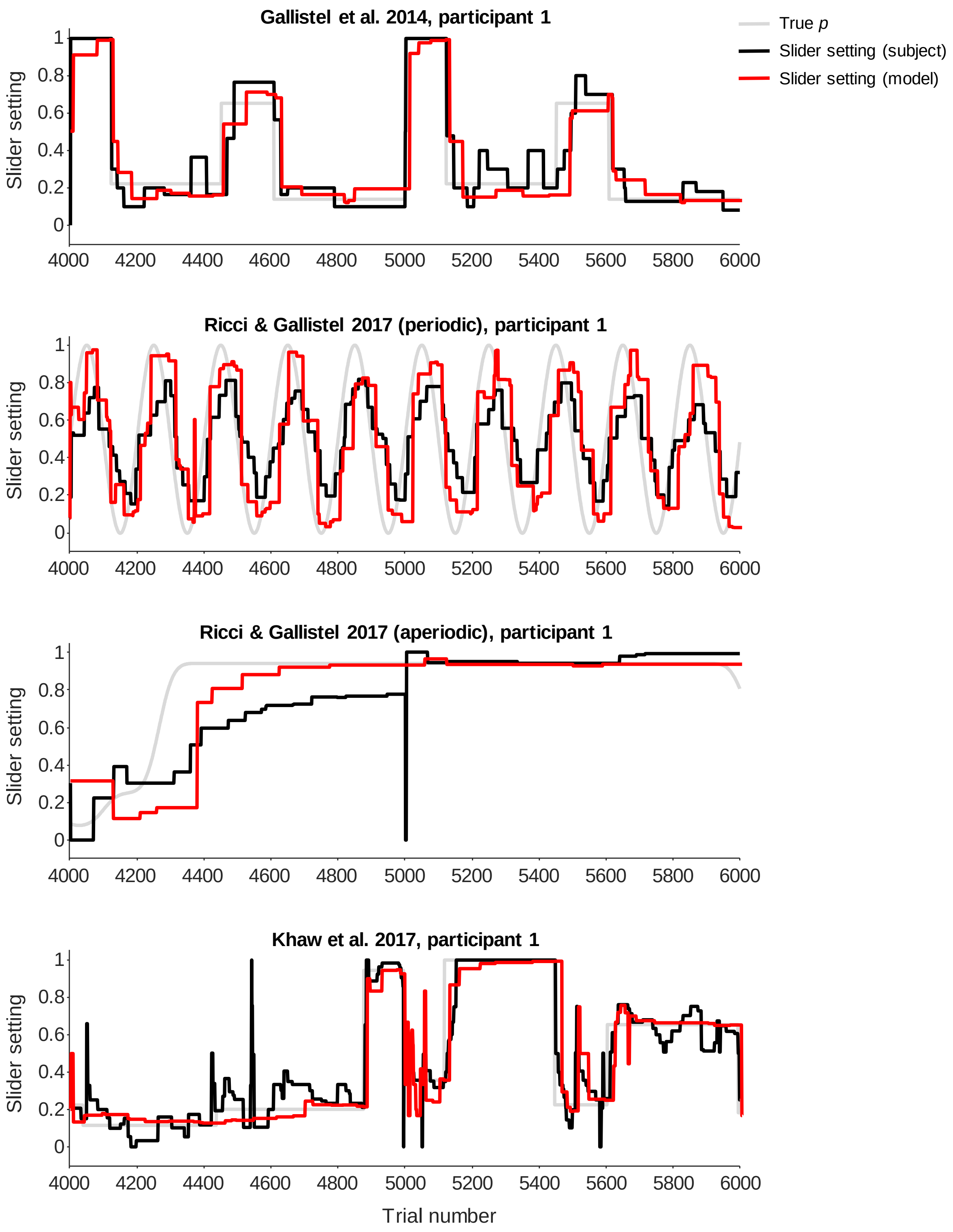


**Figure S6 | Examples of trial-by-trial slider settings of the IIAB model extended with a response threshold.** The predictions were generated by simulating responses from the model under maximum-likelihood parameter estimates. For visualisation purposes, only the central 2,000 trials are shown for each dataset.

**Robustness checks**

All model comparisons in the main paper were done with models that included a lapse rate and response noise. The lapse rate was fixed to 1/1000 in all models and we assumed Beta-distributed response noise for which the variance was fitted as a free parameter. To verify that our conclusions do not critically depend on these choices, we performed a number of additional comparisons under different assumptions about the lapse rate and response noise.

First, we compared the models under a lapse rate that is ten times larger (1/100) than in the original analysis (1/1000). The results (Figure S7A) are comparable to our main results. In particular, the delta-rule model outperformed the IIAB model by 131±20 points and was the selected model for 28 out of 29 participants, which is similar to the difference of 119±20 we found in the main analysis. Likewise, when we made the lapse rate ten times smaller (1/10000), the difference was 128±21 points and the delta-rule model was the selected model for 26 out of 29 participants (Figure S7B).

Next, we checked that the results are robust against changes in assumptions about the response noise distribution. First, we compared the models after removing response noise altogether, which produced a difference of 95±18 log likelihood points in favour of the delta-rule model, which won in the comparison for 26 out of 29 participants (Figure S7C). We also verified that the results are robust against changes in the assumed shape of the response noise distribution. To this end, we refitted the models using a truncated Gaussian instead of Beta distribution for the response noise. The results (Figure S7D) are again similar to the main results, with the delta-rule model outperforming the IIAB model on 27 out of 29 participants with a mean difference of 150±22 log likelihood points.

Finally, we verified that the model comparison results do not critically rely on using cross-validated log likelihoods as the evaluation measure. To this end, we performed an additional model comparison using the Akaike Information Criterion (Akaike, 1974) as the evaluation measure. The results are highly comparable under both measures (Figure S7E), with the delta-model being favoured for 27 out of 29 participants with an average AIC difference of 275±41 points.

These results indicate that our main model comparison results did not critically depend on assumptions made about lapse rates, response noise or on the chosen model comparison method.

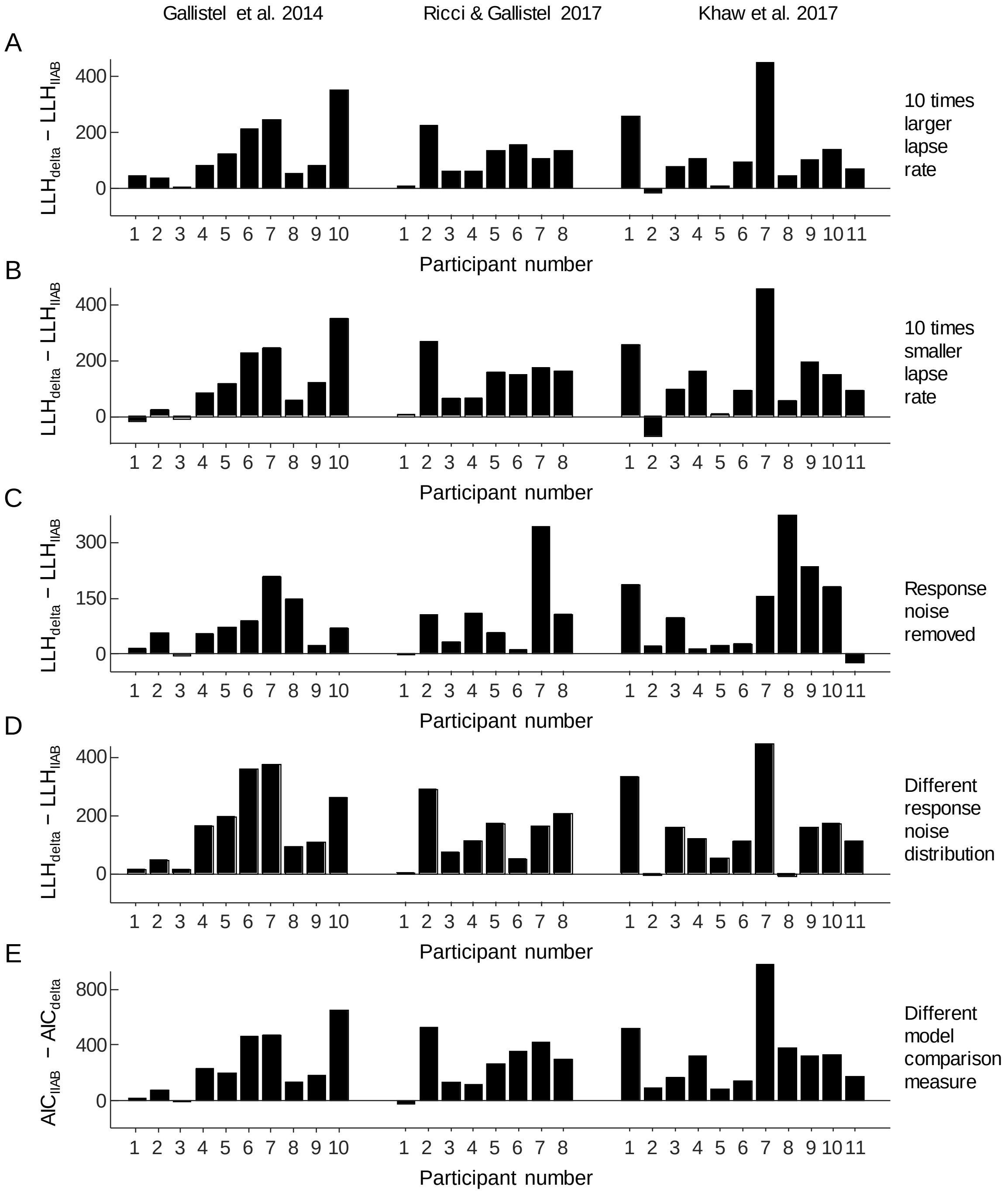


**Figure S7 | Model comparison robustness checks.** The comparisons shown in this figure are the same as the main model comparison shown in Figure 5C, except for the following differences: (A) lapse rate set to 1/100; (B) lapse rate set to 1/1000; (C) response noise removed; (D) truncated Gaussian response noise distribution; (E) models compared using AIC values instead of cross-validated likelihoods.

### Two-kernel Delta-Rule Model

Under conditions where there are large and infrequent changes, as in much of the experiment data considered in this study, the standard version of the delta-rule faces a problem. If a lot of weight is put on the most recent history (by having a high learning rate), the model will quickly catch on to changes but exhibit excessive volatility during the long periods where the true probability is unchanged. If, on the other hand, recency is given only a little weight, the model will avoid excessive volatility but be slow to catch on to sudden changes. As a potential solution, Gallistel et al. (2014) considered a two-kernel variant that keeps track of two running averages with different learning rates. The model switches between these two running averages, allowing it to keep up with sudden changes while avoiding excessive volatility. To do so, the model keeps two estimates of the Bernoulli probability, $p_{\text{slow},t} = (1-\lambda_{\text{slow}})p_{\text{slow},t-1} + \lambda_{\text{slow}}X_t$ and$p_{\text{fast},t} = (1-\lambda_{\text{fast}})p_{\text{fast},t-1} + \lambda_{\text{fast}}X_t$. On trials where the absolute difference between the two estimates is larger than a threshold $\Delta_c$, the model takes $p_{\text{fast}}$ as its estimate of the Bernoulli probability; otherwise it uses $p_{\text{slow}}$ as its estimate. The model thus has two additional parameters compared to the standard delta-rule model. We combined this updating mechanism with the same variable response threshold as was used in the standard delta model. We find that this variant outperforms the standard model by 38.8 ± 6.1 log-likelihood points. Having the flexibility to weight evidence differently at different times thus seems to increase goodness of fit sufficiently to justify the additional parameters.

### An Approximately Bayesian Delta-Rule Model

Nassar et al. (2010) suggested a delta-rule variant inspired by the same Bayesian change point detection model (Adams & MacKay, 2007) as the IIAB model. Their "approximately Bayesian delta-rule model" explicitly considers two hypotheses after each new observation: either there has been a change in the true, covert probability or there has not. Unlike the IIAB model, it performs no discrete hypothesis testing but instead balances the relative evidence of these two possibilities trial-by-trial. This balancing can be rewritten (see Nassar et al., 2010, for details) as a delta-rule with an adaptive learning rate:

$$P_{est,t} = P_{est,t-1} + \frac{1 + \Omega_t \hat{r}_t}{\hat{r}_t + 1} \times (X_t - P_{est,t})$$

$X_t$ is the latest observation and $P_{est,t}$ the estimate on trial $t$. $\Omega_t$, the probability of a change point on trial t to some unknown value, is calculated as:

$$\Omega_t = \frac{U(X_t|0,1)^{\lambda} H}{U(X_t|0,1)^{\lambda} H + B(X_t|n = 1, P_{est,t})^{\lambda}(1 - H)}$$

where $U$ and $B$ are uniform and binomial distributions[2], H is a free parameter within the range (0,1) and $\lambda$ is a parameter within the range [0,1]. When $\lambda$ is not equal to 1, the model underweights the likelihoods. $\hat{r}_{t+1}$, the "expected run length" between change points, is calculated as:

$$\hat{r}_{t+1} = (\hat{r}_t + 1)(1 - \Omega_t) + \Omega_t$$

We find that this model outperforms the IIAB model by 70 ± 25 log likelihood points, but performs worse than the regular delta-rule model by 55 ± 20 log-likelihood points.

Nassar et al. (2010) also suggested a non-normative variant that allows underweighting of likelihoods by raising them to a power. When the power is equal to 0, this model reduces to the regular delta-rule model for all but the first few trials (and can thus not perform much worse than that model). This non-normative variant performs better than the regular delta-rule model, by 31 ± 11 log-likelihood points, and does so by heavily underweighting the likelihoods (Figure S8).

In sum, this version of an adaptive learning rate does seem to improve on the regular delta-rule model if it is allowed to deviate from normativity.

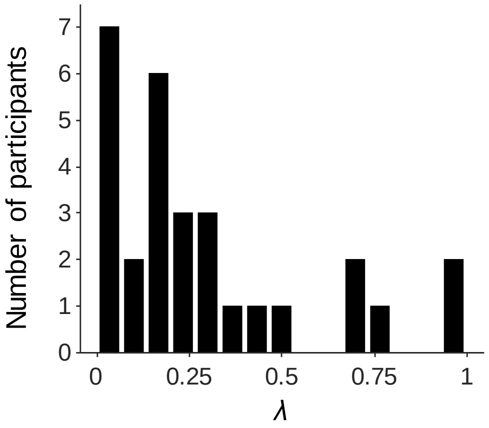


**Figure S8 |** Maximum-likelihood estimates of the likelihood weight parameter ($\lambda$) in the approximately Bayesian delta-rule model.

[2] In Nassar et al. (2010), participants draw from a Gaussian why the original version of this model uses the uniform and normal distributions. In the present experiments, participants draw from Bernoulli distributions why we use the uniform and binomial distributions.
